# Contrasting approaches to genome-wide association studies impact the detection of resistance mechanisms in *Staphylococcus aureus*

**DOI:** 10.1101/758144

**Authors:** Nicole E. Wheeler, Sandra Reuter, Claire Chewapreecha, John A. Lees, Beth Blane, Carolyne Horner, David Enoch, Nicholas M. Brown, M. Estée Török, David M. Aanensen, Julian Parkhill, Sharon J. Peacock

**Author notes:** These authors contributed equally (Nicole E. Wheeler, Sandra Reuter). Corresponding author: Nicole E. Wheeler, Centre for Genomic Pathogen Surveillance Wellcome Sanger Institute, Wellcome Genome Campus Hinxton, United Kingdom.

## Abstract

Rapid detection of antibiotic resistance using whole-genome sequencing (WGS) could improve clinical outcomes and limit the spread of resistance. For this to succeed, we need an accurate way of linking genotype to phenotype, that identifies new resistance mechanisms as they appear. To assess how close we are to this goal, we characterized antimicrobial resistance determinants in >4,000 *Staphylococcus aureus* genomes of isolates associated with bloodstream infection in the United Kingdom and Ireland. We sought to answer three questions: 1) how well did known resistance mechanisms explain phenotypic resistance in our collection, 2) how many previously identified resistance mechanisms appeared in our collection, and 3) how many of these were detectable using four contrasting genome-wide association study (GWAS) methods. Resistance prediction based on the detection of known resistance determinants was 98.8% accurate. We identified challenges in correcting for population structure, clustering orthologous genes, and identifying causal mechanisms in rare or common phenotypes, which reduced the recovery of known mechanisms. Limited sensitivity and specificity of these methods made prediction using GWAS-discovered hits alone less accurate than using literature-derived genetic determinants. However, GWAS methods identified novel mutations associated with resistance, including five mutations in *rpsJ*, which improved tetracycline resistance prediction for 28 isolates, and a T118I substitution in *fusA* which resulted in better fusidic acid resistance prediction for 5 isolates. Thus, GWAS approaches in conjunction with phenotypic testing data can support the development of comprehensive databases to enable real-time use of WGS for patient management.

## Introduction

Bacterial whole genome sequencing (WGS) is increasingly being introduced into diagnostic and public health microbiology laboratories (Raven et al. 2018). Applications beyond research include surveillance, outbreak investigation and transmission tracking (Mellmann et al. 2011; Köser et al. 2012; Gire et al. 2014), as well as the genetic prediction of antimicrobial susceptibility (Reuter et al. 2013; Walker et al. 2015; Mason et al. 2018).

*Staphylococcus aureus* was used as a model organism in some of the earliest studies that explored the clinical utility of WGS, including transmission at scales that ranged from local to global (Harris et al. 2010; Köser et al. 2012; Aanensen et al. 2016; Reuter et al. 2016; Coll et al. 2017). A case for the added value of WGS in MRSA outbreak investigations has been clearly made (Harris et al. 2013; Coll et al. 2017), and attention is now turning to additional uses for the sequence data being generated.

The current standard for determining the antimicrobial susceptibility of *S. aureus* is culture-based phenotypic testing. Culture-based results can be generated within 24 hours, but the development of sequencing technologies that produce data in a few hours could mean that genetic prediction becomes available earlier than the results of phenotypic assays. Accurate prediction depends on a robust, comprehensive database of known resistance mechanisms. Several studies have measured the accuracy of genetic prediction in *S. aureus* (Gordon et al. 2014; Bradley et al. 2015; Aanensen et al. 2016; Davis et al. 2016; Mason et al. 2018), and shown a concordance of 98-99% between genotype and phenotype measured by disc diffusion assay, in line with error from laboratory phenotypic testing (Aanensen et al. 2016). Maintaining the accuracy of genetic prediction of phenotypic resistance requires an on-going process to detect novel mechanisms as they appear (Hicks et al. 2019).

Data on resistance genes or mutations have largely been accumulated from studies using classical molecular biology techniques including gene knock-out and complementation (Read and Massey 2014). With increasing sequencing data available, this is being complemented by genome-wide association studies (GWAS) (Chen and Shapiro 2015; Falush 2016). Several bacterial GWAS have been reported for *S. aureus* (Alam et al. 2014; Laabei et al. 2014; Baines et al. 2015; Young et al. 2019) and other species (Howell et al. 2014; Earle et al. 2016; Coll et al. 2017). These have largely addressed pathogenicity and virulence (Howell et al. 2014; Laabei et al. 2014; Lees et al. 2016, 2019), and antibiotic resistance (Alam et al. 2014; Desjardins et al. 2016; Earle et al. 2016; Coll et al. 2017). They have also highlighted the challenges of bacterial GWAS, including clonal populations with little recombination that lead to strong linkage disequilibrium, and variable gene content in the accessory genome (Earle et al. 2016; Lees et al. 2016).

The strong clonality and low recombination rate within *S. aureus* can present a challenge in identifying resistance mechanisms. Because resistance is well characterised in *S. aureus*, we can test the ability of GWAS to identify known resistance mechanisms to indicate how effectively this methodology might identify novel mechanisms. Here, we describe a study of 4,140 *S. aureus* isolates, in which we characterize antimicrobial resistance determinants, confirm the accuracy of genetic resistance predictions, and examine the utility of GWAS for detecting known and putative novel genetic resistance mechanisms that will contribute to the accuracy of future databases.

## Results

### Strain collection

We analysed 4,140 *S. aureus* isolates associated with human carriage and disease in the UK and Republic of Ireland. All isolates were sequenced previously (Reuter et al. 2016; Coll et al. 2017; Donker et al. 2017), and most were resistant to 3-5 antibiotics (Fig. 1A). The collection included five dominant clonal complexes (CC22 containing the epidemic strain EMRSA-15 (n=2802), CC30 containing EMRSA-16 (n=436), CC5 (n=168), CC8 in which USA300 resides (n=147), and CC1 (n=145)), together with 13 other CCs, and sequence types (STs) that do not fall into a clonal complex (Fig. 1B). The source of isolates and method of susceptibility testing are shown in Supplemental Table S1, and further details are available in the Methods section. The population structure of our collection is heavily influenced by longitudinal MRSA sampling carried out in the East of England, and differs from the European population structure in Aanensen *et al.* (2016), with CC22 dominating our collection. The most frequent resistance profile was the co-occurrence of the four most common resistance phenotypes - penicillin, oxacillin, ciprofloxacin and erythromycin (Fig. 1C).

**Fig. 1 A:**
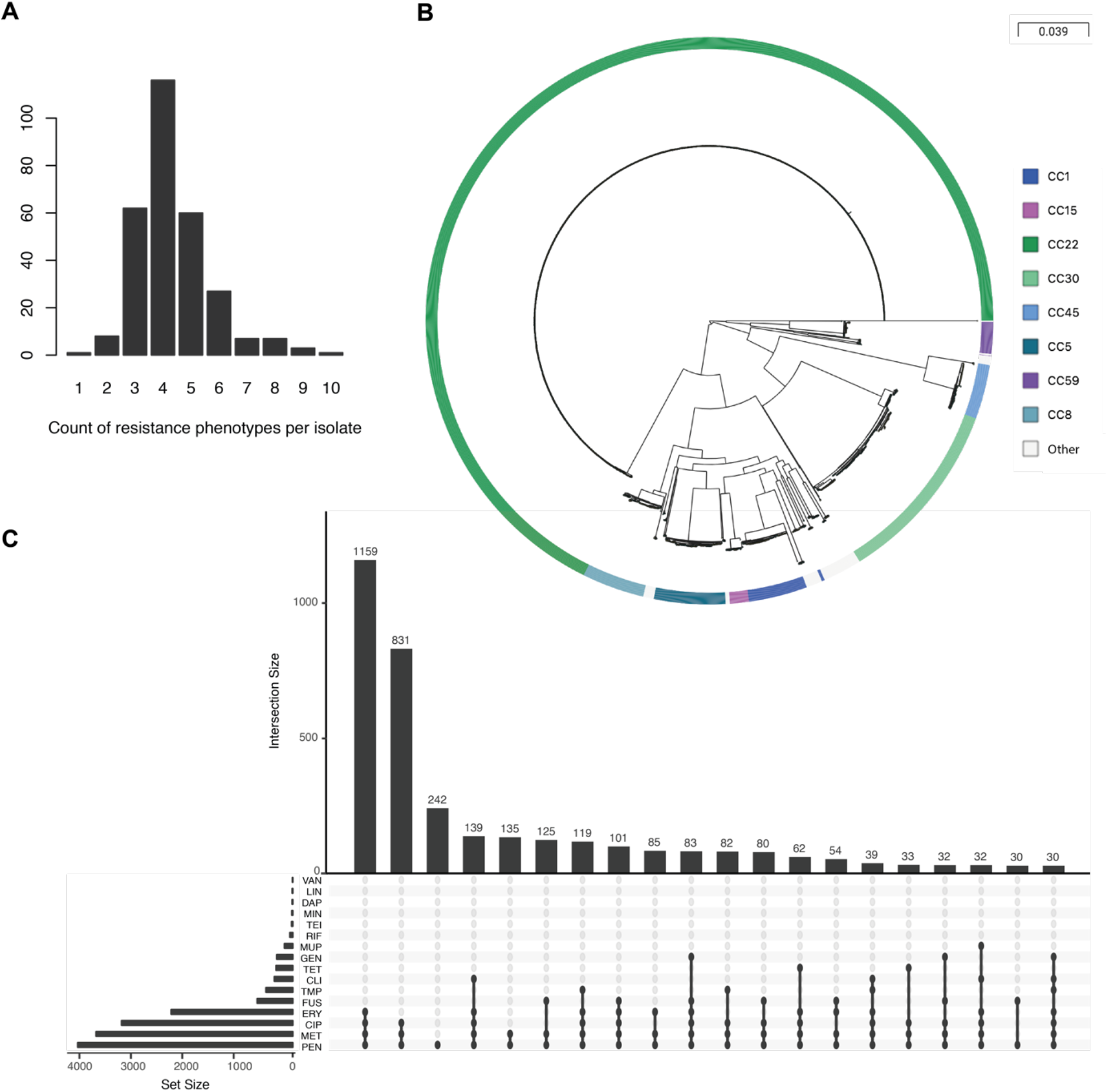
Histogram of the number of resistance phenotypes per isolate; B: Phylogenetic overview of clonal complexes in the collection. An interactive view can be accessed at: https://microreact.org/project/SaureusGWAS; C: The 20 most common resistance profiles across our collection.

### Antimicrobial resistance prediction from whole genome sequencing

We constructed a genetic resistance database, combining previous databases with additional mechanisms from the literature (see Supplemental Table S2 for details). Agreement between phenotypic antimicrobial susceptibility and genotype was 98.8% overall, and over 98% for 13/16 individual antibiotics (Fig. 2, Supplemental Table S3), consistent with previous studies (Gordon et al. 2014; Bradley et al. 2015; Aanensen et al. 2016; Mason et al. 2018). Sensitivity was over 96% for gentamicin, penicillin, oxacillin, ciprofloxacin, minocycline and trimethoprim. For erythromycin, agreement (96.5%), sensitivity (96.4%), and specificity (96.6%) were low compared with other drugs (Fig. 2). Of 145 discrepant isolates, 63 were false positives and 82 were false negatives (Supplemental Table S3). *ermC* is often present on a small plasmid in CC22, whose presence fluctuates in the population (Holden et al. 2013; Reuter et al. 2016), and in isolates from the same patient (Harris et al. 2013). We found *ermC* on a small contig in 86.3% of cases, with read coverage higher than core genes, suggesting it was located on this small plasmid. Of the discrepant isolates, 29 (all CC22) carried *ermC* but were phenotypically susceptible, and 59 isolates from CC22 were phenotypically resistant but lacked a known genetic determinant (Supplemental Table S4). Because we did not phenotype and sequence the same colony, it is possible that the plasmid was variably present within culture, or was lost during processing prior to DNA extraction for sequencing. Low sensitivity for vancomycin, teicoplanin, daptomycin and linezolid could be due to the low frequency (<1%) of resistance phenotypes, or absence of phenotypic testing data in this and other collections (Gordon et al. 2014; Mason et al. 2018).

**Fig. 2:**
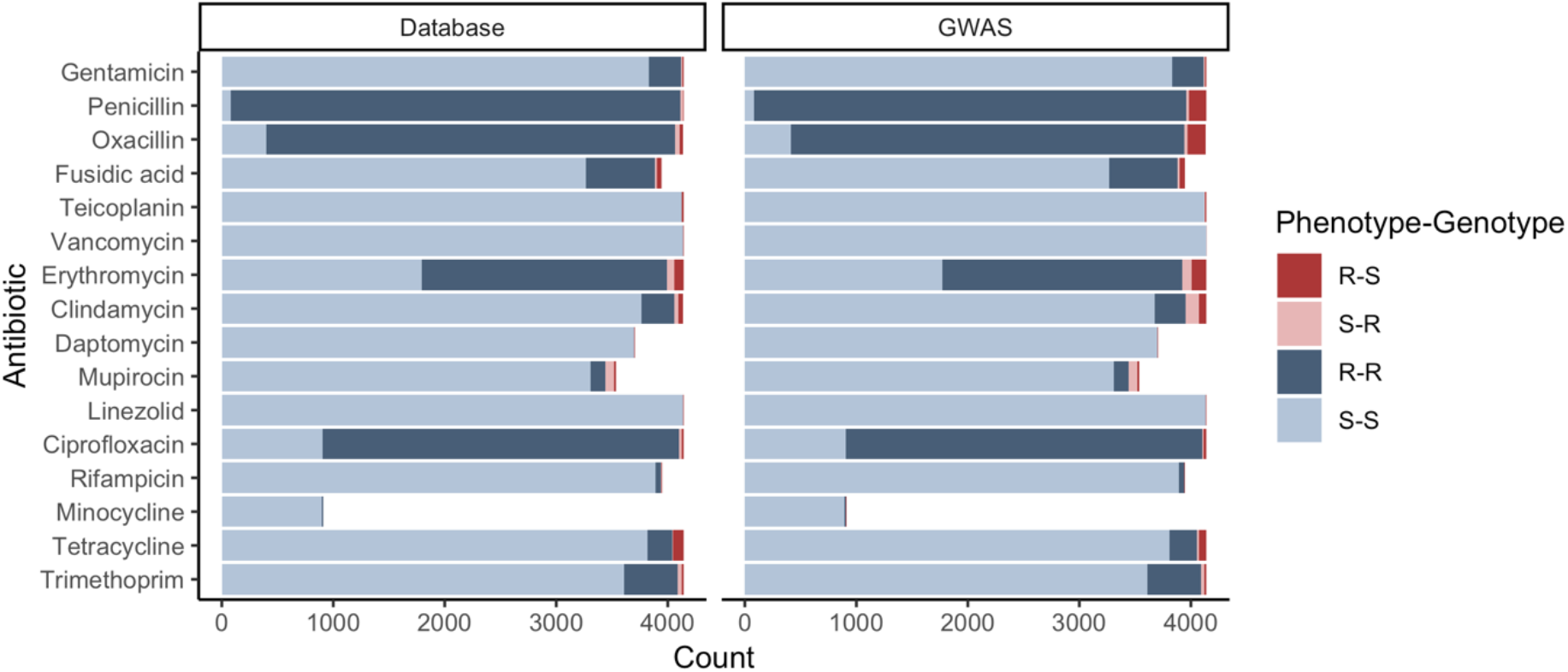
Concordance between phenotype and genotype for predictions made using a database of resistance determinants, or plausible GWAS hits. In the BSAC collection (Reuter et al. 2016), not all antibiotics were phenotyped each year, leading to fewer test results for fusidic acid, daptomycin, mupirocin, and rifampicin. The BSAC collection was the only collection for which minocycline phenotypes were available for 90% of isolates.

Specificity of genetic prediction was generally above 96%, the exceptions being the beta-lactam antibiotics penicillin (77.45%) and oxacillin (90.70%). Of 102 penicillin susceptible isolates, 18 contained intact genes with no unique mutations in their promoters or coding sequences, that encode penicillin resistance (*mecA* n=6, *blaZ* n=12). A similar observation was made by Bradley et al. (2015), and may be due to a failure of conventional phenotypic testing methods to detect penicillin resistance reliably (El Feghaly et al. 2012). Similarly, of 441 isolates that were oxacillin susceptible, 41 (9.3%) contained either intact *mecA* (n=28) or *mecC* (n=13). The 13 *mecC* isolates were phenotyped using the VITEK instrument, which can fail to detect phenotypic resistance associated with this element (Cartwright et al. 2013). Overall, 267 sensitive strains carried intact homologs of known resistance genes. This suggests that better pseudogene detection could improve specificity.

Lower concordance was also observed for mupirocin (97.3%) compared with rates of 98.9-100% described previously (Gordon et al. 2014; Bradley et al. 2015; Aanensen et al. 2016). 78 isolates were false positives, 19 were false negatives (Supplemental Table S3). Our inclusion of intermediate phenotypes may be driving this difference (Gordon et al. 2014; Bradley et al. 2015; Mason et al. 2018). Intermediate resistance is caused by mutations in *ileS-1*. The resulting minimum inhibitory concentrations (MICs) are near the epidemiological cut-off used to define resistance. Phenotypic variation of the isolate and reading variation of phenotypic tests may lead to false positives. 73/78 false positive calls were due to mutations in *ileS-1*. Of the 19 false negative calls, 14 had an intermediate phenotype, and when re-tested, 11 were susceptible (Supplemental Table S4).

### Identification of resistance loci with GWAS

We performed GWASs using four popular methods to identify resistance mechanisms. We chose methods which employ different approaches to correct for population structure. We compared FaST-LMM (Lippert et al. 2011), which uses SNP calls to compute kinship, an implementation of Pyseer (Lees et al. 2018) which uses a phylogenetic tree to measure relatedness, and treeWAS (Collins and Didelot 2018), which uses the phylogenetic tree to identify genotype-phenotype correlations that are stronger than expected based on shared ancestry and recombination. Due to the computation time required to run treeWAS, it was only applied to gene presence/absence data. We also performed a Plink analysis that corrected for clonal complex, but not finer-scale population structure. We provide code and data to compare the methods we have tested with other approaches at: https://github.com/nwheeler443/staph_gwas.

GWAS recovered known resistance mechanisms for a range of antibiotics in both the core and accessory genome (Table 1, Fig. 3), leading to good recovery of predictive ability (Fig. 2, overall accuracy of 0.981, Supplemental Table S3). To anticipate what was discoverable in our collection, we compared known resistance determinants gathered from the literature to those present in our samples. We found that for antibiotics with few genetic determinants, most were present in our dataset. However, for antibiotics with greater diversity of determinants, such as the gene variants involved in erythromycin resistance or the SNPs involved in fusidic acid resistance, fewer than half were present in our collection (Fig. 3). Of 15 known linezolid resistance SNPs and one known gene, one SNP was present in our collection, and no known determinants of resistance to teicoplanin and vancomycin were present in our samples.

**Table 1:**
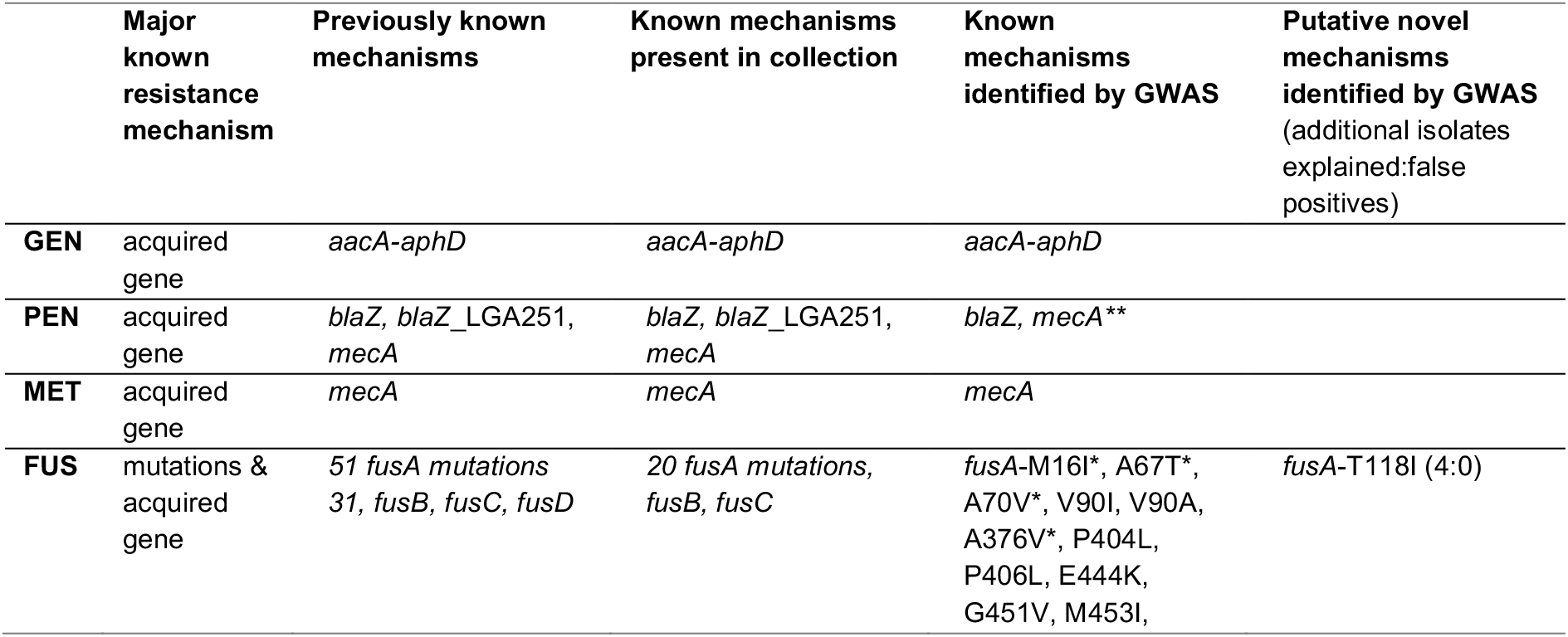
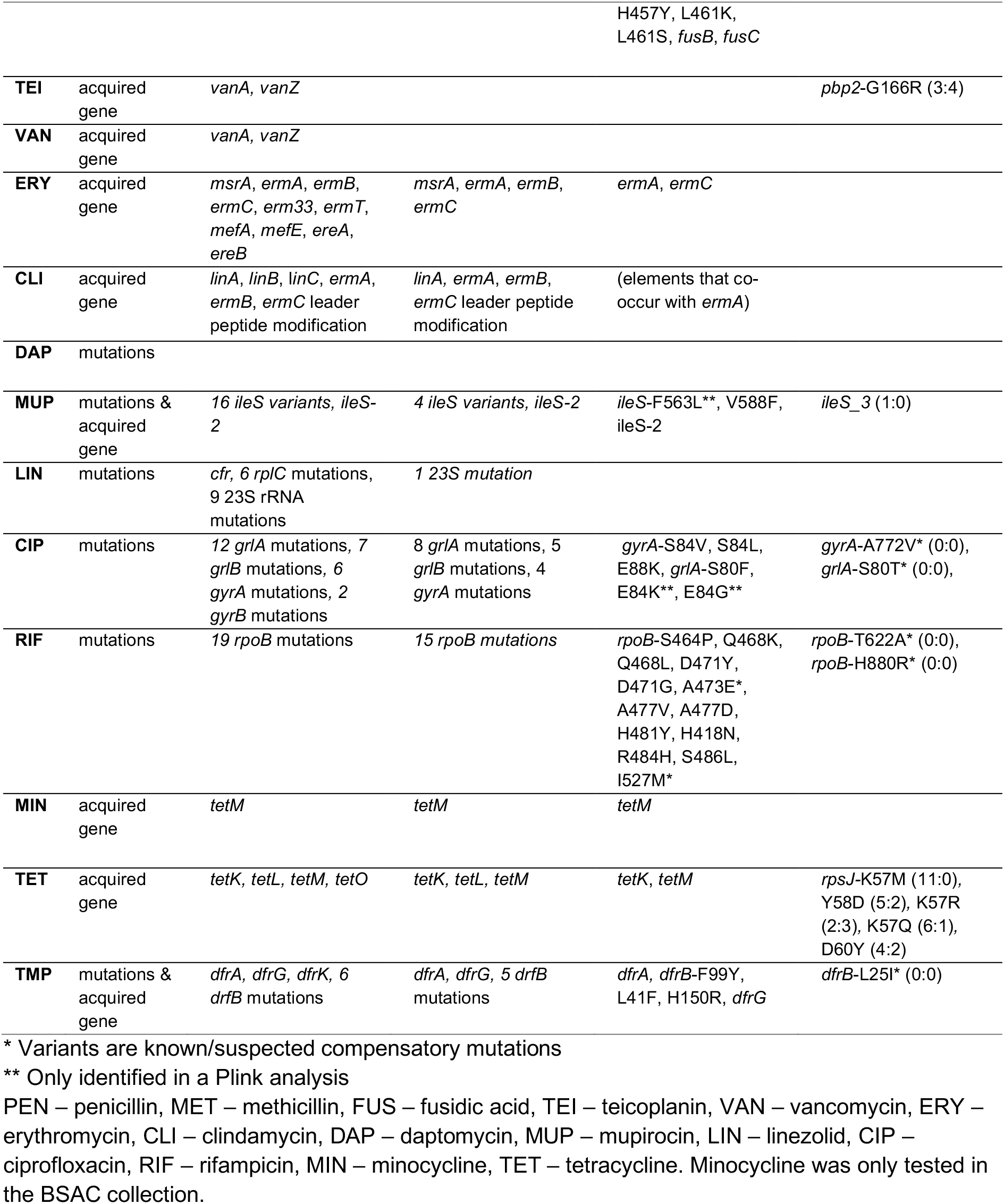
Overview of resistance mechanisms identified by at least one GWAS method. See Supplemental Table S2 for a full list of known resistance mechanisms.

|  | Major known resistance mechanism | Previously known mechanisms | Known mechanisms present in collection | Known mechanisms identified by GWAS | Putative novel mechanisms identified by GWAS (additional isolates explained:false positives) |
| --- | --- | --- | --- | --- | --- |
| **GEN** | acquired gene | *aacA-aphD* | *aacA-aphD* | *aacA-aphD* |  |
| **PEN** | acquired gene | *blaZ*, *blaZ_LGA251*, *mecA* | *blaZ*, *blaZ_LGA251*, *mecA* | *blaZ*, *mecA* ** |  |
| **MET** | acquired gene | *mecA* | *mecA* | *mecA* |  |
| **FUS** | mutations & acquired gene | 51 *fusA* mutations<br>31, *fusB*, *fusC*, *fusD* | 20 *fusA* mutations,<br>*fusB*, *fusC* | *fusA*-M16I*, A67T*, A70V*, V90I, V90A, A376V*, P404L, P406L, E444K, G451V, M453I, | *fusA*-T118I (4:0) |
|  |  |  |  | H457Y, L461K,<br>L461S, *fusB*, *fusC* |  |
| **TEI** | acquired gene | *vanA*, *vanZ* |  |  | *pbp2*-G166R (3:4) |
| **VAN** | acquired gene | *vanA*, *vanZ* |  |  |  |
| **ERY** | acquired gene | *msrA*, *ermA*, *ermB*,<br>*ermC*, *erm33*, *ermT*,<br>*mefA*, *mefE*, *ereA*,<br>*ereB* | *msrA*, *ermA*, *ermB*,<br>*ermC* | *ermA*, *ermC* |  |
| **CLI** | acquired gene | *linA*, *linB*, *linC*, *ermA*,<br>*ermB*, *ermC* leader<br>peptide modification | *linA*, *ermA*, *ermB*,<br>*ermC* leader peptide<br>modification | (elements that co-<br>occur with *ermA*) |  |
| **DAP** | mutations |  |  |  |  |
| **MUP** | mutations & acquired gene | 16 *ileS* variants, *ileS*-2 | 4 *ileS* variants, *ileS*-2 | *ileS*-F563L**, V588F,<br>*ileS*-2 | *ileS*_3 (1:0) |
| **LIN** | mutations | *cfr*, 6 *rplC* mutations,<br>9 23S rRNA<br>mutations | 1 23S mutation |  |  |
| **CIP** | mutations | 12 *griA* mutations, 7<br>*griB* mutations, 6<br>*gyrA* mutations, 2<br>*gyrB* mutations | 8 *griA* mutations, 5<br>*griB* mutations, 4<br>*gyrA* mutations | *gyrA*-S84V, S84L,<br>E88K, *griA*-S80F,<br>E84K**, E84G** | *gyrA*-A772V* (0:0),<br>*griA*-S80T* (0:0), |
| **RIF** | mutations | 19 *rpoB* mutations | 15 *rpoB* mutations | *rpoB*-S464P, Q468K,<br>Q468L, D471Y,<br>D471G, A473E*,<br>A477V, A477D,<br>H481Y, H418N,<br>R484H, S486L,<br>I527M* | *rpoB*-T622A* (0:0),<br>*rpoB*-H880R* (0:0) |
| **MIN** | acquired gene | *tetM* | *tetM* | *tetM* |  |
| **TET** | acquired gene | *tetK*, *tetL*, *tetM*, *tetO* | *tetK*, *tetL*, *tetM* | *tetK*, *tetM* | *rpsJ*-K57M (11:0),<br>Y58D (5:2), K57R<br>(2:3), K57Q (6:1),<br>D60Y (4:2) |
| **TMP** | mutations & acquired gene | *dfrA*, *dfrG*, *dfrK*, 6<br>*dfrB* mutations | *dfrA*, *dfrG*, 5 *dfrB*<br>mutations | *dfrA*, *dfrB*-F99Y,<br>L41F, H150R, *dfrG* | *dfrB*-L25I* (0:0) |

\* Variants are known/suspected compensatory mutations
\*\* Only identified in a Plink analysis
PEN – penicillin, MET – methicillin, FUS – fusidic acid, TEI – teicoplanin, VAN – vancomycin, ERY – erythromycin, CLI – clindamycin, DAP – daptomycin, MUP – mupirocin, LIN – linezolid, CIP – ciprofloxacin, RIF – rifampicin, MIN – minocycline, TET – tetracycline. Minocycline was only tested in the BSAC collection.

**Fig. 3:**
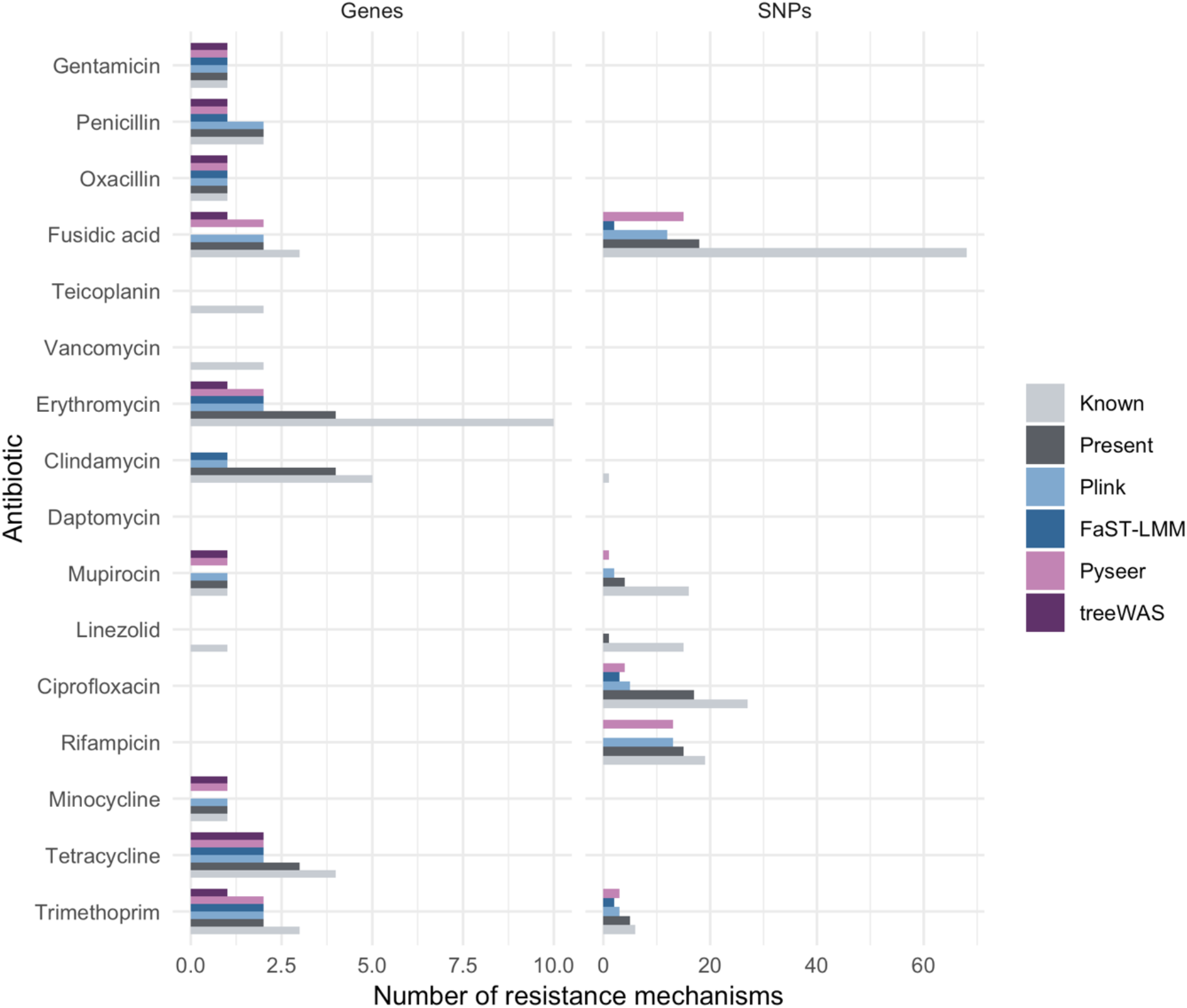
Counts of known resistance mechanisms, number that were present in our collection, and number discovered by the four GWAS methods we tested.

### GWAS recovers known resistance determinants when mechanisms are simple and phenotypes are intermediate in frequency

#### Gentamicin

The major known mechanism for gentamicin resistance in *S. aureus* is the *aacA-aphD* gene. This was identified by all four GWAS methods (Tables 1, 2, Fig. 3). Other SNPs and accessory genes showing similar patterns of presence/absence across the tree were also identified. Accounting for *aacA-aphD* explained resistance in all but nine isolates, making discovery of additional mechanisms unlikely.

**Table 2:**
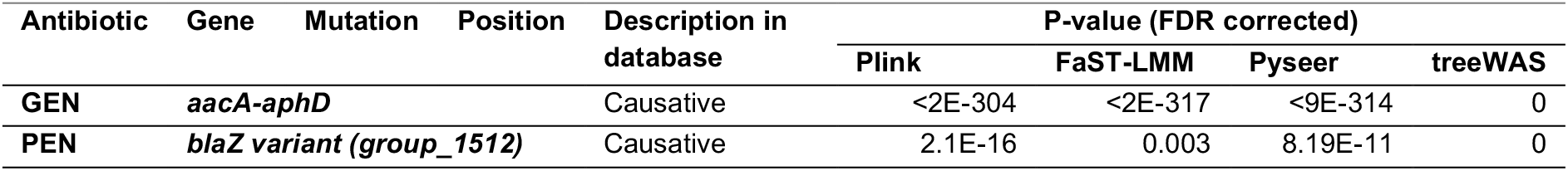
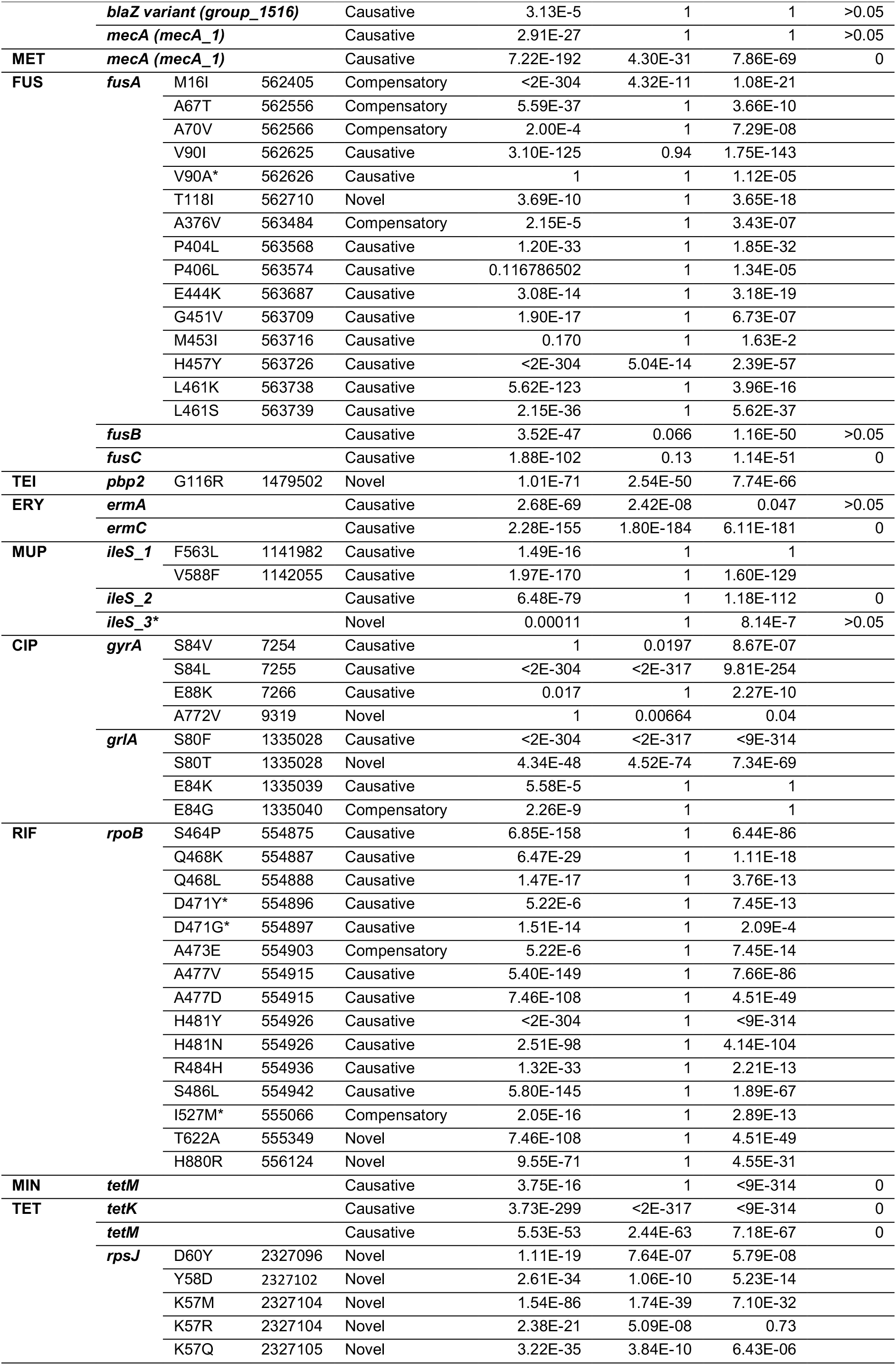
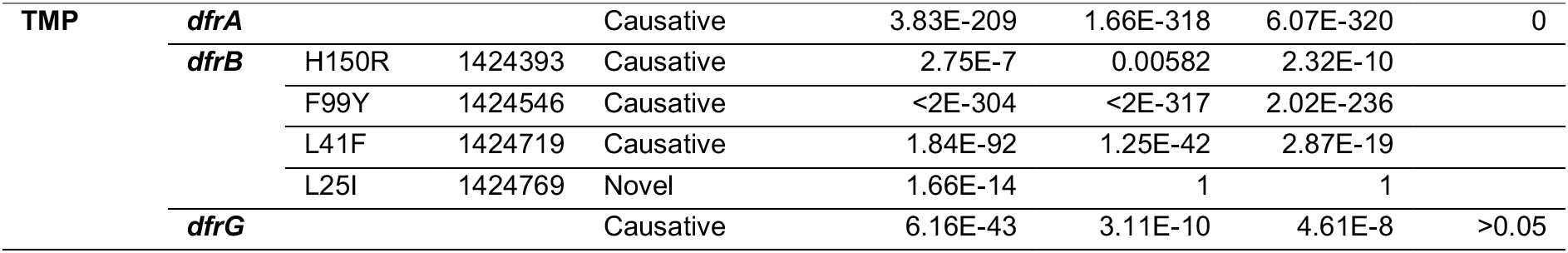
Summary of significant SNP associations. treeWAS analysis was carried out only for gene presence/absence, so P-values for SNPs are not reported. Results are reported if they appear biologically plausible and have emerged multiple times in our collection, or were present in our original database of resistance determinants. Candidates tagged with an asterisk were reported previously, but do not show homoplasy in our collection. Genome position refers to EMRSA-15 (HE681097).

#### Oxacillin

All methods successfully identified *mecA* and SCC*mec* elements as significantly associated with oxacillin resistance. Some SCC*mec* genes appeared to tag resistant strains better than *mecA*, which was due to *mecA* being split into multiple families during orthologous gene clustering.

#### Erythromycin

SNPs in *grlA* and *gyrA* were among the top associations for the FaST-LMM analysis, and manual inspection showed a clear association with phenotype. These SNPs confer resistance to ciprofloxacin, but not erythromycin. Erythromycin and ciprofloxacin resistance were strongly correlated in our collection across multiple CCs (Fisher’s exact test P-value <10^−10^), suggesting these are false positives caused by co-selection. The major plausible determinants identified were homologs of *ermA* and *ermC*, however the splitting of these genes into multiple families during gene clustering caused a loss of sensitivity in detecting these associations. 66 strains carried no known resistance determinant. 23S rRNA mutations confer resistance to erythromycin (Roberts 2008), however there were no SNPs in the 23S rRNA exclusive to resistant isolates that could explain resistance.

#### Rifampicin

Plink and Pyseer recovered 11 known causative and two known compensatory *rpoB* SNPs in an association test for rifampicin resistance (Tables 1, 2). They also identified two novel SNPs (T622A and H880R), which have been observed previously, but have always been found with other variants (Krishna 2007). In our study they each only occurred once, preventing us from confidently assigning causality. None of these SNPs were recovered by FaST-LMM.

#### Trimethoprim

GWAS methods identified three SNPs in *dfrB* previously known to confer resistance to trimethoprim. *dfrA* and *dfrG* were both identified by GWAS, as well as *thyA*, which is found in tandem with *dfrA*. Three resistant isolates were missed by our GWAS hits, due to *dfrB*-H31N or *dfrB*-L21V mutations which were too rare to be re-discovered. *dfrB*-H150R was significantly associated with resistance, but was present in more susceptible (15) than resistant/intermediate (11) isolates. A novel association identified by Plink, *dfrB*-L25I, was only seen with other known resistance mutations suggesting it could be compensatory. 15 resistant and three intermediate isolates remained unexplained.

#### Minocycline

*tetM* was identified as a resistance determinant for minocycline by Pyseer, treeWAS and Plink, but not FaST-LMM. There were no resistant isolates that did not carry *tetM*, meaning there was no additional resistance to explain.

### GWAS fails to identify resistance determinants in very rare or very common phenotypes

#### Penicillin

The major cause of penicillin resistance, the *blaZ* gene, was split into 11 subfamilies during orthologous gene clustering, with identical sequences split into different families (Supplemental Fig. S1). All GWAS methods found weak associations between one *blaZ* subfamily and penicillin resistance. Plink also identified a second *blaZ* subfamily and

#### mecA

High-frequency phenotypes are problematic for GWAS analyses, which may have also contributed to low sensitivity (The CRyPTIC Consortium and the 100,000 Genomes Project 2018). SNPs in *blaZ*, *blaR1* and the nearby *penI* were positively associated with phenotype, suggesting they acted as proxies for gene presence. Mutations in the *mecA* promoter can impact penicillin resistance, but we did not find any promising candidates d.

#### Vancomycin

Resistance to vancomycin occurred in two unrelated isolates, and there were no genes or SNPs which tagged both. The genetic basis for vancomycin intermediate and hetero-resistance (VISA and hVISA) may involve a number of regulatory and cell wall associated genes (Alam et al. 2014). A previous GWAS implicated *rpoB-*H481 mutations in the VISA phenotype (Alam et al. 2014). Several isolates carried *rpoB-*H481NYD mutations, which mostly had an MIC for vancomycin of 1-2µg/ml where MIC values were available, lower than that reported by Alam et al (2014). This can be due to methodological differences (agar dilution versus disc diffusion), as noted previously (Alam et al 2014). These MICs, and hVISA in general, are below EUCAST breakpoints, so were classified as susceptible.

#### Clindamycin

The *ermA* gene was only significantly associated with resistance in the Plink analysis. Elements of SCC*mec* and Tn*554*, the transposon that carries *ermA*, were the top hits in our Pyseer GWAS. Failure to identify *ermA* was likely due to presence of the gene in 32 sensitive strains. While inducible clindamycin resistance is conferred by *ermA*, constitutive resistance can be achieved via deletion of the *ermC* leader peptide or disruption of its position relative to the gene by an IS element (Holden et al. 2013). As the *ermC* leader peptide is only 19 amino acids long, it is not annotated by gene prediction models. Manual inspection of the leader peptide identified disruption in 33 resistant isolates.

#### Daptomycin

Seven isolates were resistant to daptomycin. No GWAS method identified a significantly associated SNP or gene common to more than one resistant strain. One isolate carried a previously identified daptomycin resistance mutation - *mprF*-L341S (Mehta et al. 2012).

#### Linezolid

Six isolates were resistant to linezolid, and none shared a plausible, significantly associated SNP or gene. Linezolid resistance can be achieved through mutation in the 23S rRNA, however because this is a multi-copy gene, mutations are often missed by haploid SNP calling. Manual examination revealed that a G2576T mutation was present on two 23S copies in a single linezolid resistant isolate.

### GWAS identifies putative novel resistance determinants

#### Fusidic acid

A strong signal for SNPs in *fusA* was identified by Plink and Pyseer, but not FaST-LMM (Fig. 4A, B). We identified 14 previously known causative or compensatory mutations, and one novel SNP in *fusA*, T118I, which explained an additional five resistant isolates from two separate emergences of resistance on the phylogenetic tree (Fig. 4C). Causal SNPs often ranked higher than compensatory SNPs (Table 2). Only two of these SNPs were identified by FaST-LMM (H57Y and M16I) (Fig. 4B, Table 2). In addition to these SNPs, *fusB* and *fusC* were tagged by Pyseer and Plink.

**Fig. 4A:**
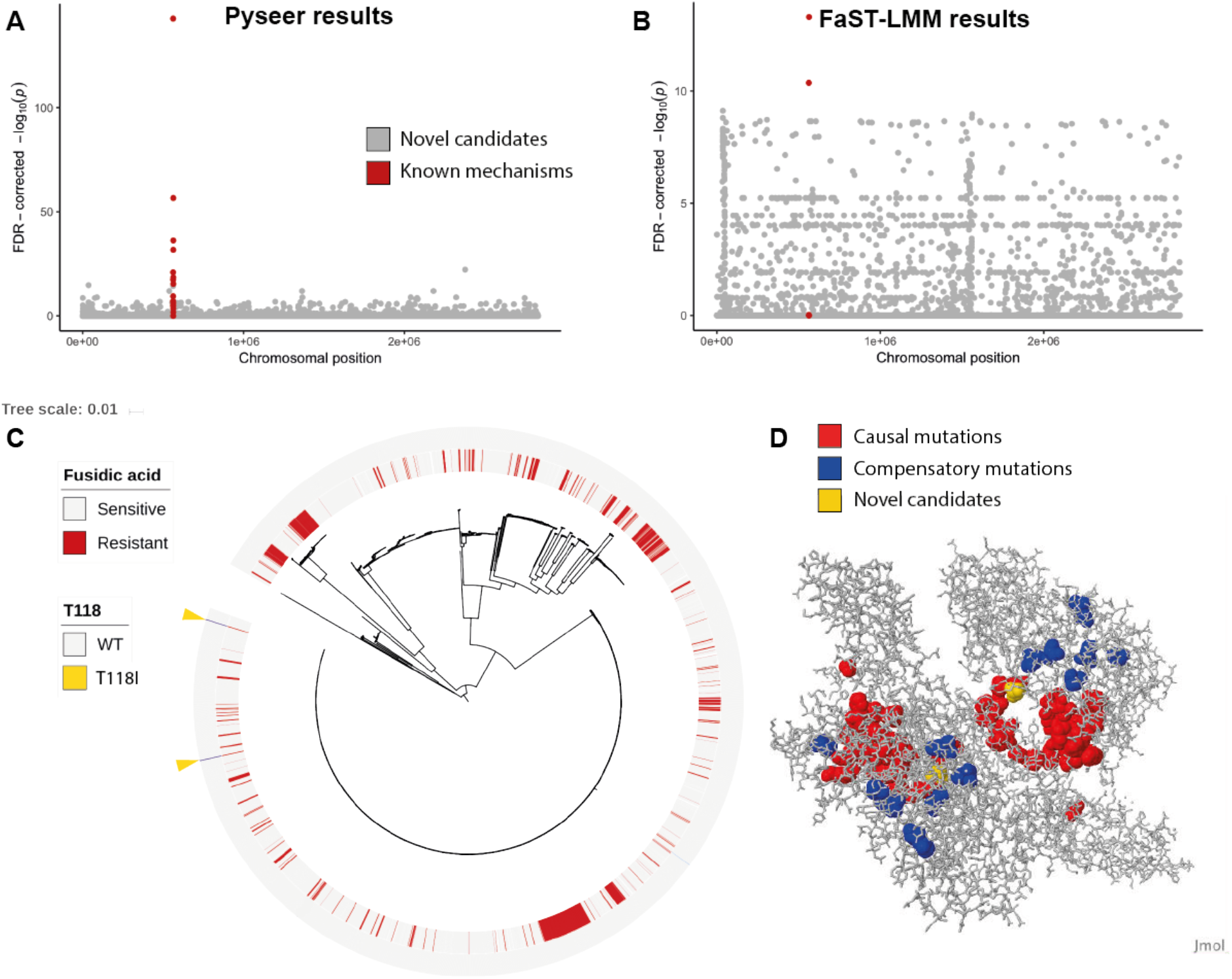
Manhattan plot for Pyseer GWAS SNP results; B: Manhattan plot for FaST-LMM GWAS SNP results, note the difference in y-axis scale; C: Newly identified GWAS SNP mapped against the *S. aureus* phylogeny; D: crystal structure 3ZZ0 of *fusA* from *Staphylococcus aureus (Koripella et al. 2012)*, with residues identified as significant in the GWAS highlighted.

The elongation factor *FusA* is the main chromosomal target for fusidic acid resistance SNPs. The novel candidate (T118I) is close to known causative mutations P114H and Q115L (Fig. 4D), and causes fusidic acid resistance in *Salmonella* Typhimurium (Johanson and Hughes 1994). We randomly down-sampled to 250, 500, 1000, 2000 and 3000 samples and re-ran a Pyseer GWAS, to assess the value of collecting more samples. As we included more samples, recovery of known mechanisms increased in a somewhat linear fashion (Supplemental Fig. S2).

#### Teicoplanin

Despite being a rare phenotype, all methods identified one SNP in *pbp2* which corresponded to three independent emergences of teicoplanin resistance. The SNP also tagged four susceptible strains, suggesting it was not sufficient to cause resistance. Teicoplanin acts by binding to peptidoglycan precursors, preventing peptidoglycan cross-linking. Increased expression of PBP2 has been observed in experimentally derived teicoplanin resistant strains (Moreira et al. 1997), however, we found no documentation of resistance mutations in this gene.

#### Mupirocin

*ileS-2* and mutations in *ileS-1* were identified by GWAS, and explained resistance well (leaving four resistant and 13 intermediate isolates unexplained). The top Plink and Pyseer SNP, *ileS-1*-V588F, was found in 91 resistant and 65 susceptible isolates. The SNP confers low level resistance, and is a first-step mutation which is followed by other mutations to increase MIC (Hurdle et al. 2004). *ileS-2* was split into multiple gene families. The gene families successfully identified two resistant isolates that had been missed by a BLAST search, and identified a variant of *ileS-2* which did not cause resistance. We also detected a novel gene, annotated as *ileS-3*. This gene was similar to *ileS-2* (78.61% identity; Supplemental Fig. S3). While it was only present in one isolate, it achieved statistical significance in the Pyseer and Plink analyses (Table 2).

#### Ciprofloxacin

*gyrA*-S84L and *grlA*-S80F explained 99% of ciprofloxacin resistance, and were identified by all GWAS methods. 10 of 3,164 isolates with *gyrA*-S84L were susceptible to ciprofloxacin. All methods identified *grlA*-S80T, which was often (59/60 isolates) found with the *gyrA*-S84L SNP, suggesting it was compensatory. A subset of approaches (Table 2) also identified three SNPs in *gyrA* (known causative mutations S84V and E88K, and novel SNP A772V, which was found with known causative SNPs so may be compensatory), and two mutations in *grlA* (E84K, a known causal mutation and E84G, a known compensatory mutation).

#### Tetracycline

*tetK* was the top association across all four methods, followed by *tetM*, collectively accounting for 90% of fully resistant isolates. Gene clustering split *tetM* into three families, one which was significantly associated with resistance across all four methods and one that was significantly associated for all methods except treeWAS. 91 isolates had no acquired resistance genes but had intermediate (68) or full (23) resistance to tetracycline. Tetracycline acts by binding to the small subunit of the ribosome and interfering with aminoacyl tRNA docking (Wilson 2014). All three tested GWAS methods highlighted *rpsJ*, which encodes the S10 ribosomal protein of the small 30S ribosomal subunit, as a target for mutations that could explain intermediate resistance.

Amino acids 55-58 of *rpsJ* were the most variable in our collection (Fig. 5A). Consistent with previous observations (Beabout et al. 2015), the most common site for significant associations was K57 (Fig. 5B). The mutations significantly associated with resistance (*rpsJ*-K57MQR, Y58D and D60Y) have been observed in *S. aureus* or *E. faecium* in a previous experimental evolution study which selected for resistance to tigecycline. Allelic replacement to verify these mutations have failed to produce a viable mutant (Beabout et al. 2015; Cattoir et al. 2015). Studies in *Neisseria gonorrhoeae* have also described a V57M intermediate resistance mutation in the corresponding site (Hu et al. 2005). These mutation hotspots overlap in *S. aureus*, *Thermus thermophilus*, and *N. gonorrhoeae* (Hu et al. 2005), (Fig. 5A) and are close to the antibiotic binding site (Brodersen et al. 2000; Hu et al. 2005) (Fig. 5C). Including these mutations in prediction improved sensitivity from 71% to 80%.

**Fig. 5A:**
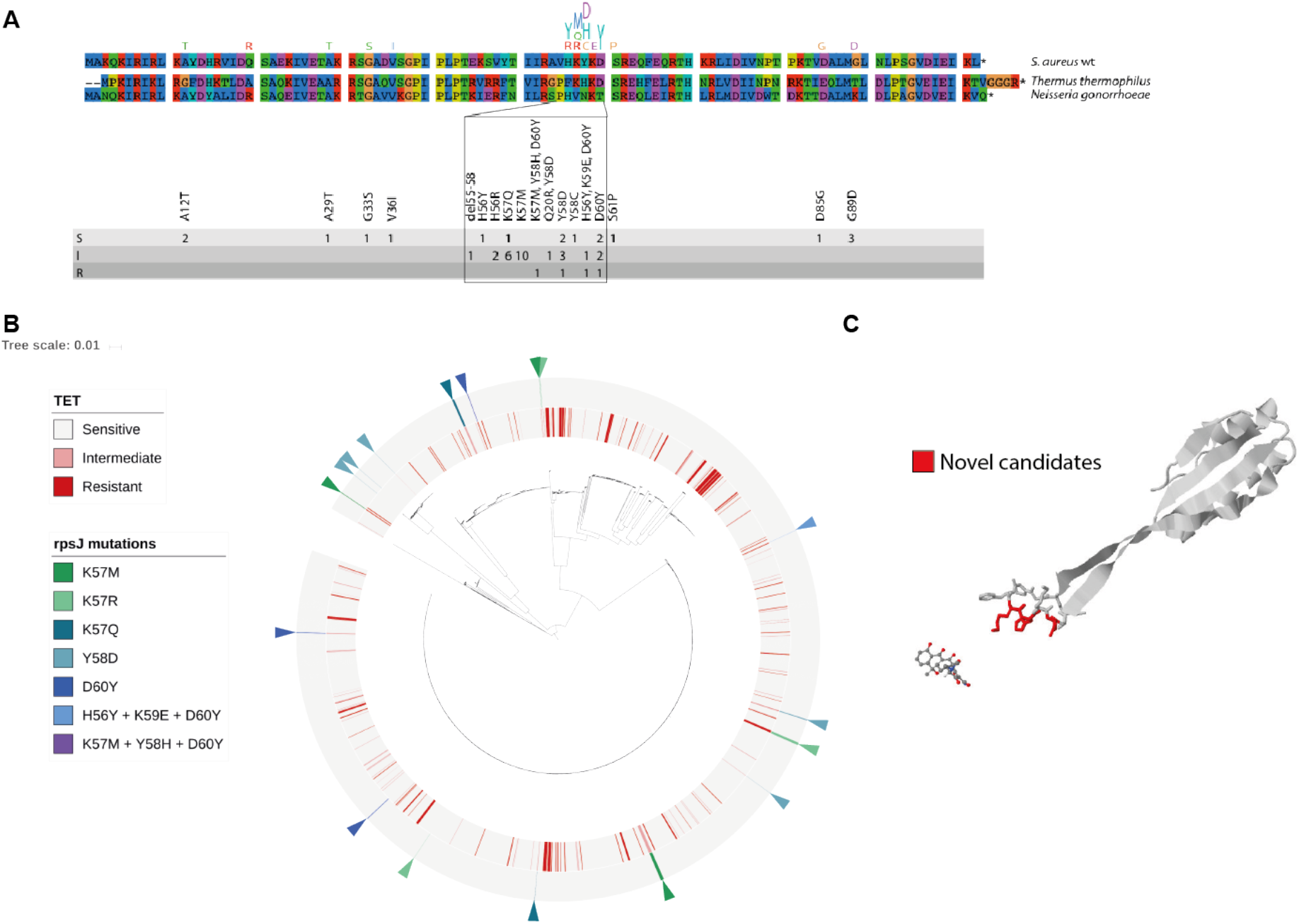
Alignment of RpsJ from *S. aureus, T. Thermophilus* and *N. gonorrhoeae*, indicating mutations found in our collection and their correspondence to phenotype (S,I,R); B: GWAS SNPs mapped against the *S. aureus* phylogeny; C: significant GWAS residues in RpsJ from *Thermus thermophilus* crystal structure 4V9A (Jenner et al. 2013), with tetracycline shown in its binding site.

### Accuracy of resistance prediction using GWAS hits

We used Pyseer GWAS results to test the ability of GWAS hits to predict phenotype, as it provided a good trade-off in sensitivity and specificity. First, we classified strains as resistant if they carried SNPs or genes with significant, positive associations with resistance. This approach predicted all isolates in our study to be pan-resistant, reflecting the high burden of false positive associations identified by GWAS (Tam et al. 2019). We also tested stricter P-value (P=10^−5^ and 10^−10^) and beta cutoffs (0.1 and 0.2), however these did not result in high accuracy (median accuracy: 12%, IQR: 1-55%). Next, we employed the elastic net function in Pyseer to train a model for predicting resistance. We partitioned the collection into 18 clades using fastbaps (Tonkin-Hill et al. 2019) to re-weight the samples for elastic net fitting and split the strains into folds for cross-validation. The performance of this method was high, but did not reach the level achieved by curation of the literature (Table 3). This approach performed the worst when predicting fusidic acid and erythromycin resistance, where a number of gene variants contribute to resistance. However, manual curation of the GWAS hits found across the four methods using prior knowledge of resistance mechanisms resulted in accuracy of 98.1% (Fig.2).

**Table 3:** Performance of elastic net models for predicting resistance compared to the performance of a database of resistance determinants curated from the literature.

| Antibiotic | True positives | True negatives | False positives | False negatives | Elastic net accuracy | Accuracy of database |
| --- | --- | --- | --- | --- | --- | --- |
| GEN | 7 | 3471 | 0 | 226 | 94% | 99% |
| PEN | 3606 | 0 | 98 | 0 | 97% | 99% |
| MET | 3327 | 270 | 94 | 9 | 97% | 98% |
| FUS | 191 | 2901 | 9 | 451 | 87% | 98% |
| TEI | 0 | 3693 | 0 | 11 | 100% | 100% |
| VAN | 0 | 3702 | 0 | 2 | 100% | 100% |
| ERY | 1774 | 618 | 1122 | 190 | 66% | 97% |
| CLI | 7 | 3604 | 0 | 93 | 97% | 98% |
| DAP | 2 | 3336 | 0 | 4 | 100% | 100% |
| MUP | 2 | 3125 | 0 | 84 | 97% | 97% |
| LIN | 0 | 3698 | 0 | 6 | 100% | 100% |
| CIP | 2802 | 713 | 69 | 120 | 95% | 99% |
| RIF | 1 | 3510 | 0 | 41 | 99% | 100% |
| MIN | - | - | - | - | - | 100% |
| TET | 0 | 3417 | 0 | 287 | 92% | 98% |
| TMP | 9 | 3303 | 0 | 392 | 89% | 99% |

### Splitting of gene families during clustering hampered association testing

Known resistance gene families (*aacA-aphD*, *blaZ*, *mecA*, *ermC*, *ermA*, and *ileS-2*) were broken into several families, some of which were not significantly associated with phenotype. However, clustering of *ile-2* was beneficial in splitting BLAST hits into variants that correlated well with resistance and those that didn’t (Supplemental Table S6). Clustering of orthologous genes differs depending on the sequence identify cutoff used to separate gene families. Our cutoff of 90% was chosen to balance splitting and merging families, but the best cutoff differed for each gene.

### Population structure correction using a phylogenetic tree is more effective than using SNP calls

We identified many significantly associated SNPs and genes that could not be plausibly linked to resistance. The majority of these are likely to be false positives. Method of correction for population structure had a strong effect on our ability to filter true associations from false (Fig. 4, Table 2). Pyseer and treeWAS often ranked known mechanisms higher than FaST-LMM (Table 2). When FaST-LMM and Pyseer are both run using SNP calls to compute kinship, they show similar difficulty excluding lineage effects. treeWAS was more conservative in reporting significant hits. The Plink analysis recovered more causal mechanisms for penicillin, oxacillin, fusidic acid and mupirocin resistance, but reported vastly more significant associations overall (Supplemental Figure S4).

### Manual examination of plausible candidates provides a complement to GWAS

Due to the clear concentration of resistance mutations in some proteins, we performed a manual candidate gene search to complement the GWAS. All mutations identified with this approach occurred in only one isolate, making their direct detection by GWAS impossible. We noted one isolate with *fusA*-E468A (MIC 2µg/ml), one with E468V (8µg/ml), and one with E468K, A70V and novel substitution V299A (MIC 128µg/ml). Given the proximity of other causal mutations, these are promising candidates. We found 15 isolates with H457C, a new substitution at a known position. We identified the known resistance mutation *grlA*-E84V in 11 ciprofloxacin resistant isolates, and *grlA*-E84Q in three related resistant isolates. *rpsJ*-H56R, previously observed in a tigecycline resistant *E. faecium* strain (Bender et al. 2018), was also found in two isolates exhibiting intermediate tetracycline resistance, but sensitive to vancomycin and teicoplanin. We observed variants in the large ribosomal unit L4 that may be associated with erythromycin resistance. 18 isolates carried mutations *rplD*-G69A and T70P, which have been described previously in two sporadic clinical *S. aureus* isolates as causative for erythromycin resistance (Prunier et al. 2005) and fall within a conserved region of the protein, known to interact with erythromycin, where similar mutations have been described in *E. coli* (Zaman et al. 2007).

## Discussion

Automated detection of resistance determinants from genomic surveillance data would substantially advance our understanding of resistance and ability to detect it. We found that in a collection of over 4,000 clinical isolates from the UK, phenotypes with simple genetic mechanisms could be explained by GWAS, while more complex mechanisms, or over- or under-represented phenotypes presented substantial challenges for GWAS approaches. This is unfortunate, given that GWAS approaches are primarily used to gain a better understanding of more complex phenotypes that are not currently well explained. In our study, we found that correction for population structure using SNP calls was inadequate for some phenotypes, while correction for population structure using a tree produced a better ratio of true to false positives. This is likely to be due to strong clonal population structure in *S. aureus*.

Novel resistance SNPs are likely to be so rare in the population that large sample sizes are needed to identify them. In our collection, previously known resistance determinants were able to explain 98.8% of resistance phenotypes, which is in line with the error rate of routine laboratory testing (Aanensen et al. 2016), but could still lead to treatment failures and enrichment for missing resistance mechanisms if these databases are used at scale to lead clinical decision making (Hicks et al. 2019). Thus, explaining as much of the remaining one percent remains an important goal. In our identification of plausible novel mechanisms using GWAS, we were able to increase accuracy to 98.9%. We found in our analysis that true resistance mechanisms are often buried among false positives, indicating that scrutiny of results and supporting experimental evidence are still crucial.

Novel mechanisms fell into three categories. The first is mutations leading to novel amino acid changes in known resistance genes, such as *fusA* for fusidic acid or *gyrA*/*grlA* for ciprofloxacin. The second captures putative novel mutations in credible candidate loci, such as *rpsJ* for tetracycline resistance and *pbp2* for teicoplanin resistance. These require experimental validation to confirm cause and effect, but their appearance in plausible genes, in independent lineages and species supports their value as candidates. The final category is novel genes and variants that may cause resistance, such as a novel *ileS* variant for mupirocin resistance.

Rare phenotypes and variants are underpowered in association studies (Read and Massey 2014; Lees et al. 2016). This is the case in vancomycin, and teicoplanin intermediate and hetero-resistance, which are rare and may involve complex mechanisms (Alam et al. 2014). Further challenges result from limitations in current approaches to data processing. Resistance arising from mutation in some but not all copies of 16S and 23S rRNA genes and mutations that affect the regulatory control of the *ermC* leader peptide are more difficult to detect than simple presence or absence of a gene or amino acid change. We also found difficulty in reconciling gene family groupings produced by automated gene clustering, which often split known resistance determinants into multiple families, weakening their association with phenotype and often strengthening their association with specific lineages. The use of unitigs (Jaillard et al. 2018) and read-mapping approaches (Hunt et al. 2017) can address these shortcomings, but interpretation of results is less clear-cut.

For some phenotypes there is a clear concentration of resistance mutations in a gene, and a clear spatial concentration of causal mutations within a protein, providing regions monitor for novel mutations associated with resistance. A similar approach was taken by Casali et al. (2014) to identify compensatory mutations associated with rifampicin resistance. Previously undiscovered resistance mechanisms are likely to be rare in the population, so flagging novel mutations in these regions for further investigation could be a powerful complement for identifying resistance mechanisms.

Antibiotic resistance prediction based on WGS data is being considered in diagnostic laboratories as methods become sufficiently robust and reliable for clinical use. An ongoing process will be needed to add new mechanisms to resistance determinant databases, which will require centralization and accreditation. We have shown that putative antibiotic resistance determinants can be identified using GWAS, homology with known resistance genes, and the concentration of causal mutations within genes. This can provide a powerful approach to prioritize candidate antimicrobial resistance loci for experimental validation.

## Materials and Methods

### Isolate collection and susceptibility testing

4,140 *S. aureus* isolates (3,736 MRSA, 404 MSSA) were included in this study. 1,011 MRSA bacteraemia isolates that were submitted to the British Society for Antimicrobial Chemotherapy (BSAC) bacteraemia resistance surveillance programme (Reynolds et al. 2008) between 2001-2010 by 47 laboratories in the United Kingdom and the Republic of Ireland (Reuter et al. 2016). A further 538 blood culture isolates (134 MRSA and 404 MSSA) were collected from patients at the Cambridge University Hospitals NHS Foundation Trust (CUH), in Cambridge, United Kingdom between 2006 and 2012, and 269 MRSA isolates were collected from 9 hospitals in the East of England between 1998 and 2012, 242 of which were from a single hospital (Donker et al. 2017). Isolates sampled in both the BSAC and regional collections were de-duplicated. 2,322 MRSA isolates associated with carriage and disease were sourced from four hospitals and 75 primary care practices in Cambridgeshire between April 2012 and April 2013 (Coll et al. 2017). 2,804 samples came from individual patients, 483 patients contributed multiple isolates (total of 1,336 isolates).

Phenotypic antimicrobial susceptibility testing of the BSAC collection was carried out by BSAC agar dilution for 16 antibiotics (ciprofloxacin, clindamycin, daptomycin, erythromycin, fusidic acid, gentamicin, linezolid, minocycline, mupirocin, oxacillin, penicillin, rifampicin, teicoplanin, tetracycline, trimethoprim, vancomycin). Susceptibility testing was not carried out for fusidic acid, mupirocin, and rifampicin in all 10 years of the collection. Antimicrobial susceptibility of the remaining isolates was determined using the VITEK 2 instrument (bioMerieux, Marcy l’Etoile, France) to all antibiotics except minocycline. Antimicrobial susceptibility results were interpreted using the MIC breakpoints agreed upon by the European Committee on Antimicrobial Susceptibility Testing (EUCAST) criteria (http://www.eucast.org/clinical_breakpoints/).

### Whole genome sequencing

DNA was extracted using the QIAxtractor instrument (QIAgen, Hilden, Germany). Index-tagged libraries were created, and 96 indexed libraries were sequenced in each of eight channels using the Illumina HiSeq platform (Illumina Inc.) to generate 75 or 100 base pair (bp) paired-end reads at the Wellcome Sanger Institute. Average coverage of 142X was achieved with average Phred scores of 37. Annotated assemblies were produced using the pipeline described in Page et al. (Page et al. 2016a). For each sample, sequence reads were used to create multiple assemblies using VelvetOptimiser v2.2.5 (Gladman et al. 2008) and Velvet v1.2 (Zerbino and Birney 2008). An assembly improvement step was applied to the assembly with the best N50 and contigs were scaffolded using SSPACE (Boetzer et al. 2011) and sequence gaps filled using GapFiller (Boetzer and Pirovano 2012). The mean number of contigs was 38. Automated annotation was performed using PROKKA v1.5 (Seemann 2014) and a genus specific databases from RefSeq (Pruitt et al. 2012).

For each sample, reads were mapped against the CC22 reference genome HO 5096 0412 (accession no. HE681097) (Köser et al. 2012) using SMALT v0.7.4 (Wellcome Sanger Institute) to produce a BAM file. SMALT was used to index the reference using a kmer size of 13 and a step size of 6 and reads were aligned using default parameters but with the maximum insert size (i) set as 3 times the mean fragment size of the sequencing library. PCR duplicate reads were identified using Picard v1.92 (Broad Institute) and flagged as duplicates in the BAM file. Variant detection was performed using samtools mpileup v0.1.19 (Li et al. 2009) with parameters “-d 1000 -DSugBf” and bcftools v0.1.19 (http://samtools.github.io/bcftools/) to produce a BCF file of all variant sites. Genotypes at variant sites were called using bcftools. The variant quality score was required to be greater than 50 (quality < 50) and mapping quality greater than 30 (map_quality < 30). If all reads did not give the same base call, the allele frequency, as calculated by bcftools, was required to be either 0 for bases called the same as the reference, or 1 for bases called as a single nucleotide polymorphism (SNP) (af1 < 0.95). The majority base call was required to be present in at least 75% of reads mapping at the base, (ratio < 0.75), and the minimum mapping depth required was 4 reads, at least two of which had to map to each strand (depth < 4, depth_strand < 2). Finally, strand_bias was required to be less than 0.001, map_bias less than 0.001 and tail_bias less than 0.001. If any of these filters were not met, the base was called as uncertain and masked. Regions containing mobile genetic elements, repetitive regions or phage were excluded. Multi-allelic sites were split using bcftools. The software developed by Pathogen Informatics at the Wellcome Sanger Institute is freely available for download from GitHub (Wellcome Sanger Institute, http://github.com/sanger-pathogens/vr-codebase) under an open source GNU GPL 3 license.

### Phylogeny construction

A pseudo-genome was constructed by substituting the base call at each site in the BCF file into the reference genome and any site called as uncertain was substituted with an N. Insertions with respect to the reference genome were ignored and deletions were filled with N. Variant sites were extracted from an alignment of pseudo-genomes using snp-sites (Page et al. 2016b). Phylogenetic estimation was then performed on this SNP alignment using RAxML with the general time reversible model and gamma correction (Stamatakis 2014).

### Accessory genome clustering

The pan-genome was determined using Roary (Page et al. 2015), using a 90% ID cutoff, as we found the default 95% cutoff to be too conservative for this collection. This resulted in a common core gene set of 1,517 genes present in 99% of isolates, and 10,934 genes present in <99% of isolates, which were used for further analysis.

### Antibiotic resistance prediction

We expanded a database containing mechanisms that confer antibiotic resistance in *S. aureus* from previous publications (Holden et al. 2013; Gordon et al. 2014; Bradley et al. 2015; Aanensen et al. 2016) to include additional resistance mechanisms gathered from the literature (Supplemental Table S2). Acquired resistance genes were detected using a BLAST search against *de novo* assemblies of each isolate (Camacho et al. 2009). Reads were mapped against a multifasta file of susceptible reference alleles to identify known SNPs encoding resistance. The genes were also extracted from the annotated assemblies and compared to susceptible alleles. 85 isolates had *ermC* read coverage lower than that of housekeeping genes, and were assigned as *ermC* negative.

To investigate indels upstream of the *ermA* gene, reads were mapped to the Tn*554* transposon using SMALT with GATK indel realignment (McKenna et al. 2010). Indels were then plotted over the regulatory region of the gene to identify deletions of inverted repeats 5 and 6. To identify disruptions in the *ermC* leader peptide, reads were mapped to *ermC*. Paired reads where only one mate-pair mapped within the gene were identified. Those would be reads that map either directly upstream or downstream of the *ermC* gene. These reads were then mapped to the region directly upstream of the *ermC* gene, which includes the leader peptide. The distribution of the mapping reads could then be to identify deletion of the leader peptide and insertion of IS elements.

### Genome-wide association studies

Association tests were performed using FaST-LMM (v.0.2.32) (Lippert et al. 2011), Pyseer (v.1.2.0) (Lees et al. 2018), and treeWAS (v.1.0) (Collins and Didelot 2018). For FaST-LMM analyses, a kinship matrix was calculated using SNP calls. For the Pyseer analysis, the kinship matrix was calculated using the phylogenetic tree. For the treeWAS analysis, we only included results from the “simultaneous” test. An association test that corrected for clonal complex was performed using the Cochran-Mantel-Haenszel test implemented in Plink (Purcell et al. 2007), using the --mh --within --adjust flags, as in (Chewapreecha et al. 2014).

### Prediction of resistance using GWAS results

We defined isolates as resistant if they carried any of the significantly positively associated SNPs or genes identified in a Pyseer GWAS. Our selection criteria were FDR-adjusted P-value<0.05, 1×10^−5^, 1×10^−10^, and beta value cutoffs of 0, 0.1 and 0.2. Both parameters were tested in all combinations. We also used Pyseer to fit an elastic net to predict resistance using SNP and gene data, using the default value for alpha (0.0069) which gives behaviour closer to ridge regression but still allows predictors to be removed. We clustered our strains using fastbaps (Tonkin-Hill et al. 2019) and the SNP alignment used to generate our phylogenetic tree. We used the clusters to re-weight the samples for elastic net construction and generate folds for cross-validation.

### Visualisation of results

Significantly associated SNPs were visualised against the phylogenetic tree of the collection using iTol (Letunic and Bork 2019). Protein structures were retrieved from the RCSB Protein Data Bank (Berman et al. 2000), and visualised in Jmol (Herráez 2006). Sequence alignments were viewed and coloured in Jalview (Waterhouse et al. 2009).

### DATA ACCESS

All sequences have been deposited in the European Nucleotide Archive, with individual accession numbers and phenotypes provided in Supplemental Table S4. The data used to perform the GWAS analysis, and the code used to produce Fig. 3 are available at https://github.com/nwheeler443/staph_gwas.

## Acknowledgments

We thank members of the East of England Microbiology Research Network for contribution of isolates and data: Benny Cherian (formerly of Basildon and Thurrock University Hospitals, Basildon, UK, now at Barts Health NHS Trust), Tony Elston (formerly of Colchester Hospital University, Colchester, UK, now retired), Richard Kent (formerly of Ipswich Hospital, Ipswich, UK, now retired), John Hayward (formerly of James Paget University Hospital, Great Yarmouth, UK, now retired), Louise Teare (Mid Essex Hospital Services Trust, Chelmsford, UK), Juliet Foweraker (lately of Royal Papworth Hospital, Papworth Everard, UK), David Enoch (formerly of Peterborough and Stamford Hospitals, Peterborough, UK, now at Public Health England Clinical Microbiology and Public Health Laboratory, Cambridge, UK), Nada Elhag (Southend University Hospital, Westcliff-on-Sea, UK), Prema Singh (West Hertfordshire Hospitals, Watford, UK), and Rebecca Tilley (West Suffolk Hospital, Bury St Edmunds, UK). We are grateful for assistance from the library construction, sequencing and core informatics teams, and Simon Harris at the Wellcome Trust Sanger Institute. The study was supported by grants from the UKCRC Translational Infection Research (TIR) Initiative, the Medical Research Council (Grant Number G1000803) with contributions to the Grant from the Biotechnology and Biological Sciences Research Council, the National Institute for Health Research on behalf of the Department of Health, the National Institutes of Health (grant number U01CA207167) supporting NEW, a Chief Scientist Office of the Scottish Government Health Directorate to SJP; a Sir Henry Wellcome Postdoctoral fellowship (Grant Number 107376//Z/15/Z) to Dr Chewapreecha; an MRC studentship (Grant Number 1365620) to JAL; a Hospital Infection Society Major Research Grant, and Wellcome Trust grant number 098051 awarded to the Wellcome Trust Sanger Institute. MET is a Clinician Scientist Fellow funded by the Academy of Medical Sciences and the Health Foundation, and supported by the NIHR Cambridge Biomedical Research Centre. The contents of this publication are solely the responsibility of the authors and do not necessarily represent the official views of the National Institutes of Health.

## References

Aanensen DM, Feil EJ, Holden MTG, Dordel J, Yeats CA, Fedosejev A, Goater R, Castillo-Ramírez S, Corander J, Colijn C, et al. 2016. Whole-Genome Sequencing for Routine Pathogen Surveillance in Public Health: a Population Snapshot of Invasive Staphylococcus aureus in Europe. MBio 7. http://dx.doi.org/10.1128/mBio.00444-16.

Alam MT, Petit RA 3rd, Crispell EK, Thornton TA, Conneely KN, Jiang Y, Satola SW, Read TD. 2014. Dissecting vancomycin-intermediate resistance in staphylococcus aureus using genome-wide association. Genome Biol Evol 6: 1174–1185.

Baines SL, Holt KE, Schultz MB, Seemann T, Howden BO, Jensen SO, van Hal SJ, Coombs GW, Firth N, Powell DR, et al. 2015. Convergent adaptation in the dominant global hospital clone ST239 of methicillin-resistant Staphylococcus aureus. MBio 6: e00080.

Beabout K, Hammerstrom TG, Perez AM, Magalhães BF, Prater AG, Clements TP, Arias CA, Saxer G, Shamoo Y. 2015. The ribosomal S10 protein is a general target for decreased tigecycline susceptibility. Antimicrob Agents Chemother 59: 5561–5566.

Bender JK, Cattoir V, Hegstad K, Sadowy E, Coque TM, Westh H, Hammerum AM, Schaffer K, Burns K, Murchan S, et al. 2018. Update on prevalence and mechanisms of resistance to linezolid, tigecycline and daptomycin in enterococci in Europe: Towards a common nomenclature. Drug Resist Updat 40: 25–39.

Berman HM, Westbrook J, Feng Z, Gilliland G, Bhat TN, Weissig H, Shindyalov IN, Bourne PE. 2000. The Protein Data Bank. Nucleic Acids Res 28: 235–242.

Boetzer M, Henkel CV, Jansen HJ, Butler D, Pirovano W. 2011. Scaffolding pre-assembled contigs using SSPACE. Bioinformatics 27: 578–579.

Boetzer M, Pirovano W. 2012. Toward almost closed genomes with GapFiller. Genome Biol 13: R56.

Bradley P, Gordon NC, Walker TM, Dunn L, Heys S, Huang B, Earle S, Pankhurst LJ, Anson L, de Cesare M, et al. 2015. Rapid antibiotic-resistance predictions from genome sequence data for Staphylococcus aureus and Mycobacterium tuberculosis. Nat Commun 6: 10063.

Broad Institute. Picard: A set of command line tools (in Java) for manipulating high-throughput sequencing (HTS) data. https://broadinstitute.github.io/picard/.

Brodersen DE, Clemons WM Jr, Carter AP, Morgan-Warren RJ, Wimberly BT, Ramakrishnan V. 2000. The structural basis for the action of the antibiotics tetracycline, pactamycin, and hygromycin B on the 30S ribosomal subunit. Cell 103: 1143–1154.

Camacho C, Coulouris G, Avagyan V, Ma N, Papadopoulos J, Bealer K, Madden TL. 2009. BLAST+: architecture and applications. BMC Bioinformatics 10: 421.

Cartwright EJP, Paterson GK, Raven KE, Harrison EM, Gouliouris T, Kearns A, Pichon B, Edwards G, Skov RL, Larsen AR, et al. 2013. Use of Vitek 2 antimicrobial susceptibility profile to identify mecC in methicillin-resistant Staphylococcus aureus. J Clin Microbiol 51: 2732–2734.

Casali N, Nikolayevskyy V, Balabanova Y, Harris SR, Ignatyeva O, Kontsevaya I, Corander J, Bryant J, Parkhill J, Nejentsev S, et al. 2014. Evolution and transmission of drug-resistant tuberculosis in a Russian population. Nat Genet 46: 279–286.

Cattoir V, Isnard C, Cosquer T, Odhiambo A, Bucquet F, Guérin F, Giard J-C. 2015. Genomic analysis of reduced susceptibility to tigecycline in Enterococcus faecium. Antimicrob Agents Chemother 59: 239–244.

Chen PE, Shapiro BJ. 2015. The advent of genome-wide association studies for bacteria. Curr Opin Microbiol 25: 17–24.

Chewapreecha C, Marttinen P, Croucher NJ, Salter SJ, Harris SR, Mather AE, Hanage WP, Goldblatt D, Nosten FH, Turner C, et al. 2014. Comprehensive identification of single nucleotide polymorphisms associated with beta-lactam resistance within pneumococcal mosaic genes. PLoS Genet 10: e1004547.

Coll F, Harrison EM, Toleman MS, Reuter S, Raven KE, Blane B, Palmer B, Kappeler ARM, Brown NM, Török ME, et al. 2017. Longitudinal genomic surveillance of MRSA in the UK reveals transmission patterns in hospitals and the community. Sci Transl Med 9. http://dx.doi.org/10.1126/scitranslmed.aak9745.

Collins C, Didelot X. 2018. A phylogenetic method to perform genome-wide association studies in microbes that accounts for population structure and recombination. PLoS Comput Biol 14: e1005958.

Davis JJ, Boisvert S, Brettin T, Kenyon RW, Mao C, Olson R, Overbeek R, Santerre J, Shukla M, Wattam AR, et al. 2016. Antimicrobial Resistance Prediction in PATRIC and RAST. Sci Rep 6: 27930.

Desjardins CA, Cohen KA, Munsamy V, Abeel T, Maharaj K, Walker BJ, Shea TP, Almeida DV, Manson AL, Salazar A, et al. 2016. Genomic and functional analyses of Mycobacterium tuberculosis strains implicate ald in D-cycloserine resistance. Nat Genet 48: 544–551.

Donker T, Reuter S, Scriberras J, Reynolds R, Brown NM, Török ME, James R, Network EOEMR, Aanensen DM, Bentley SD, et al. 2017. Population genetic structuring of methicillin-resistant Staphylococcus aureus clone EMRSA-15 within UK reflects patient referral patterns. Microb Genom 3: e000113.

Earle SG, Wu C-H, Charlesworth J, Stoesser N, Gordon NC, Walker TM, Spencer CCA, Iqbal Z, Clifton DA, Hopkins KL, et al. 2016. Identifying lineage effects when controlling for population structure improves power in bacterial association studies. Nat Microbiol 1: 16041.

El Feghaly RE, Stamm JE, Fritz SA, Burnham C-AD. 2012. Presence of the blaZ beta-lactamase gene in isolates of Staphylococcus aureus that appear penicillin susceptible by conventional phenotypic methods. Diagn Microbiol Infect Dis 74: 388–393.

Falush D. 2016. Bacterial genomics: Microbial GWAS coming of age. Nat Microbiol 1: 16059.

Gire SK, Goba A, Andersen KG, Sealfon RSG, Park DJ, Kanneh L, Jalloh S, Momoh M, Fullah M, Dudas G, et al. 2014. Genomic surveillance elucidates Ebola virus origin and transmission during the 2014 outbreak. Science 345: 1369–1372.

Gladman S, Seemann T, Victorian Bioinformatics Consortium. 2008. Velvet Optimiser: For automatically optimising the primary parameter options for the Velvet de novo sequence assembler. http://bioinformatics.net.au/software.velvetoptimiser.shtml.

Gordon NC, Price JR, Cole K, Everitt R, Morgan M, Finney J, Kearns AM, Pichon B, Young B, Wilson DJ, et al. 2014. Prediction of Staphylococcus aureus antimicrobial resistance by whole-genome sequencing. J Clin Microbiol 52: 1182–1191.

Harris SR, Cartwright EJP, Török ME, Holden MTG, Brown NM, Ogilvy-Stuart AL, Ellington MJ, Quail MA, Bentley SD, Parkhill J, et al. 2013. Whole-genome sequencing for analysis of an outbreak of meticillin-resistant Staphylococcus aureus: a descriptive study. Lancet Infect Dis 13: 130–136.

Harris SR, Feil EJ, Holden MTG, Quail MA, Nickerson EK, Chantratita N, Gardete S, Tavares A, Day N, Lindsay JA, et al. 2010. Evolution of MRSA during hospital transmission and intercontinental spread. Science 327: 469–474.

Herráez A. 2006. Biomolecules in the computer: Jmol to the rescue. Biochem Mol Biol Educ 34: 255–261.

Hicks AL, Kissler SM, Lipsitch M, Grad YH. 2019. Quantifying the surveillance required to sustain genetic marker-based antibiotic resistance diagnostics. bioRxiv 699918. https://www.biorxiv.org/content/10.1101/699918v1?rss=1 (Accessed July 19, 2019).

Holden MTG, Hsu L-Y, Kurt K, Weinert LA, Mather AE, Harris SR, Strommenger B, Layer F, Witte W, de Lencastre H, et al. 2013. A genomic portrait of the emergence, evolution, and global spread of a methicillin-resistant Staphylococcus aureus pandemic. Genome Res 23: 653–664.

Howell KJ, Weinert LA, Chaudhuri RR, Luan S-L, Peters SE, Corander J, Harris D, Angen Ø, Aragon V, Bensaid A, et al. 2014. The use of genome wide association methods to investigate pathogenicity, population structure and serovar in Haemophilus parasuis. BMC Genomics 15: 1179.

Hu M, Nandi S, Davies C, Nicholas RA. 2005. High-level chromosomally mediated tetracycline resistance in Neisseria gonorrhoeae results from a point mutation in the rpsJ gene encoding ribosomal protein S10 in combination with the mtrR and penB resistance determinants. Antimicrob Agents Chemother 49: 4327–4334.

Hunt M, Mather AE, Sánchez-Busó L, Page AJ, Parkhill J, Keane JA, Harris SR. 2017. ARIBA: rapid antimicrobial resistance genotyping directly from sequencing reads. Microb Genom 3: e000131.

Hurdle JG, O’Neill AJ, Ingham E, Fishwick C, Chopra I. 2004. Analysis of mupirocin resistance and fitness in Staphylococcus aureus by molecular genetic and structural modeling techniques. Antimicrob Agents Chemother 48: 4366–4376.

Jaillard M, Lima L, Tournoud M, Mahé P, van Belkum A, Lacroix V, Jacob L. 2018. A fast and agnostic method for bacterial genome-wide association studies: Bridging the gap between k-mers and genetic events. PLoS Genet 14: e1007758.

Jenner L, Starosta AL, Terry DS, Mikolajka A, Filonava L, Yusupov M, Blanchard SC, Wilson DN, Yusupova G. 2013. Structural basis for potent inhibitory activity of the antibiotic tigecycline during protein synthesis. Proc Natl Acad Sci U S A 110: 3812–3816.

Johanson U, Hughes D. 1994. Fusidic acid-resistant mutants define three regions in elongation factor G of Salmonella typhimurium. Gene 143: 55–59.

Koripella RK, Chen Y, Peisker K, Koh CS, Selmer M, Sanyal S. 2012. Mechanism of elongation factor-G-mediated fusidic acid resistance and fitness compensation in Staphylococcus aureus. J Biol Chem 287: 30257–30267.

Köser CU, Ellington MJ, Cartwright EJP, Gillespie SH, Brown NM, Farrington M, Holden MTG, Dougan G, Bentley SD, Parkhill J, et al. 2012. Routine use of microbial whole genome sequencing in diagnostic and public health microbiology. PLoS Pathog 8: e1002824.

Krishna A. 2007. Functional analysis of a pleiotropic transcription regulator in Staphylococcus aureus: Rsp. Doctor of Philosophy, Imperial College London.

Laabei M, Recker M, Rudkin JK, Aldeljawi M, Gulay Z, Sloan TJ, Williams P, Endres JL, Bayles KW, Fey PD, et al. 2014. Predicting the virulence of MRSA from its genome sequence. Genome Res 24: 839–849.

Lees JA, Ferwerda B, Kremer PHC, Wheeler NE, Serón MV, Croucher NJ, Gladstone RA, Bootsma HJ, Rots NY, Wijmega-Monsuur AJ, et al. 2019. Joint sequencing of human and pathogen genomes reveals the genetics of pneumococcal meningitis. Nat Commun 10: 2176.

Lees JA, Galardini M, Bentley SD, Weiser JN, Corander J. 2018. pyseer: a comprehensive tool for microbial pangenome-wide association studies. Bioinformatics 34: 4310–4312.

Lees JA, Vehkala M, Välimäki N, Harris SR, Chewapreecha C, Croucher NJ, Marttinen P, Davies MR, Steer AC, Tong SYC, et al. 2016. Sequence element enrichment analysis to determine the genetic basis of bacterial phenotypes. Nat Commun 7: 12797.

Letunic I, Bork P. 2019. Interactive Tree Of Life (iTOL) v4: recent updates and new developments. Nucleic Acids Res. http://dx.doi.org/10.1093/nar/gkz239.

Li H, Handsaker B, Wysoker A, Fennell T, Ruan J, Homer N, Marth G, Abecasis G, Durbin R, 1000 Genome Project Data Processing Subgroup. 2009. The Sequence Alignment/Map format and SAMtools. Bioinformatics 25: 2078–2079.

Lippert C, Listgarten J, Liu Y, Kadie CM, Davidson RI, Heckerman D. 2011. FaST linear mixed models for genome-wide association studies. Nat Methods 8: 833–835.

Mason A, Foster D, Bradley P, Golubchik T, Doumith M, Gordon NC, Pichon B, Iqbal Z, Staves P, Crook D, et al. 2018. Accuracy of Different Bioinformatics Methods in Detecting Antibiotic Resistance and Virulence Factors from Staphylococcus aureus Whole-Genome Sequences. J Clin Microbiol 56. http://dx.doi.org/10.1128/JCM.01815-17.

McKenna A, Hanna M, Banks E, Sivachenko A, Cibulskis K, Kernytsky A, Garimella K, Altshuler D, Gabriel S, Daly M, et al. 2010. The Genome Analysis Toolkit: a MapReduce framework for analyzing next-generation DNA sequencing data. Genome Res 20: 1297–1303.

Mehta S, Cuirolo AX, Plata KB, Riosa S, Silverman JA, Rubio A, Rosato RR, Rosato AE. 2012. VraSR two-component regulatory system contributes to mprF-mediated decreased susceptibility to daptomycin in in vivo-selected clinical strains of methicillin-resistant Staphylococcus aureus. Antimicrob Agents Chemother 56: 92–102.

Mellmann A, Harmsen D, Cummings CA, Zentz EB, Leopold SR, Rico A, Prior K, Szczepanowski R, Ji Y, Zhang W, et al. 2011. Prospective genomic characterization of the German enterohemorrhagic Escherichia coli O104:H4 outbreak by rapid next generation sequencing technology. PLoS One 6: e22751.

Moreira B, Boyle-Vavra S, deJonge BL, Daum RS. 1997. Increased production of penicillin-binding protein 2, increased detection of other penicillin-binding proteins, and decreased coagulase activity associated with glycopeptide resistance in Staphylococcus aureus. Antimicrob Agents Chemother 41: 1788–1793.

Page AJ, Cummins CA, Hunt M, Wong VK, Reuter S, Holden MTG, Fookes M, Falush D, Keane JA, Parkhill J. 2015. Roary: rapid large-scale prokaryote pan genome analysis. Bioinformatics 31: 3691–3693.

Page AJ, De Silva N, Hunt M, Quail MA, Parkhill J, Harris SR, Otto TD, Keane JA. 2016a. Robust high-throughput prokaryote de novo assembly and improvement pipeline for Illumina data. Microb Genom 2: e000083.

Page AJ, Taylor B, Delaney AJ, Soares J, Seemann T, Keane JA, Harris SR. 2016b. SNP-sites: rapid efficient extraction of SNPs from multi-FASTA alignments. Microb Genom 2: e000056.

Pruitt KD, Tatusova T, Brown GR, Maglott DR. 2012. NCBI Reference Sequences (RefSeq): current status, new features and genome annotation policy. Nucleic Acids Res 40: D130–5.

Prunier A-L, Trong HN, Tande D, Segond C, Leclercq R. 2005. Mutation of L4 ribosomal protein conferring unusual macrolide resistance in two independent clinical isolates of Staphylococcus aureus. Microb Drug Resist 11: 18–20.

Purcell S, Neale B, Todd-Brown K, Thomas L, Ferreira MAR, Bender D, Maller J, Sklar P, de Bakker PIW, Daly MJ, et al. 2007. PLINK: a tool set for whole-genome association and population-based linkage analyses. Am J Hum Genet 81: 559–575.

Raven K, Blane B, Churcher C, Parkhill J, Peacock SJ. 2018. Are commercial providers a viable option for clinical bacterial sequencing? Microb Genom. http://dx.doi.org/10.1099/mgen.0.000173.

Read TD, Massey RC. 2014. Characterizing the genetic basis of bacterial phenotypes using genome-wide association studies: a new direction for bacteriology. Genome Med 6: 109.

Reuter S, Ellington MJ, Cartwright EJP, Köser CU, Török ME, Gouliouris T, Harris SR, Brown NM, Holden MTG, Quail M, et al. 2013. Rapid bacterial whole-genome sequencing to enhance diagnostic and public health microbiology. JAMA Intern Med 173: 1397–1404.

Reuter S, Török ME, Holden MTG, Reynolds R, Raven KE, Blane B, Donker T, Bentley SD, Aanensen DM, Grundmann H, et al. 2016. Building a genomic framework for prospective MRSA surveillance in the United Kingdom and the Republic of Ireland. Genome Res 26: 263–270.

Reynolds R, Hope R, Williams L, BSAC Working Parties on Resistance Surveillance. 2008. Survey, laboratory and statistical methods for the BSAC Resistance Surveillance Programmes. J Antimicrob Chemother 62 Suppl 2: ii15–28.

Roberts MC. 2008. Update on macrolide–lincosamide–streptogramin, ketolide, and oxazolidinone resistance genes. FEMS Microbiol Lett 282: 147–159.

Seemann T. 2014. Prokka: rapid prokaryotic genome annotation. Bioinformatics 30: 2068–2069.

Stamatakis A. 2014. RAxML version 8: a tool for phylogenetic analysis and post-analysis of large phylogenies. Bioinformatics 30: 1312–1313.

Tam V, Patel N, Turcotte M, Bossé Y, Paré G, Meyre D. 2019. Benefits and limitations of genome-wide association studies. Nat Rev Genet. http://dx.doi.org/10.1038/s41576-019-0127-1.

The CRyPTIC Consortium and the 100,000 Genomes Project. 2018. Prediction of Susceptibility to First-Line Tuberculosis Drugs by DNA Sequencing. N Engl J Med 379: 1403–1415.

Tonkin-Hill G, Lees JA, Bentley SD, Frost SDW, Corander J. 2019. Fast hierarchical Bayesian analysis of population structure. Nucleic Acids Res 47: 5539–5549.

Walker TM, Kohl TA, Omar SV, Hedge J, Del Ojo Elias C, Bradley P, Iqbal Z, Feuerriegel S, Niehaus KE, Wilson DJ, et al. 2015. Whole-genome sequencing for prediction of Mycobacterium tuberculosis drug susceptibility and resistance: a retrospective cohort study. Lancet Infect Dis 15: 1193–1202.

Waterhouse AM, Procter JB, Martin DMA, Clamp M, Barton GJ. 2009. Jalview Version 2—a multiple sequence alignment editor and analysis workbench. Bioinformatics 25: 1189–1191.

Wellcome Sanger Institute. Bio-Assembly-Improvement: Improvement of genome assemblies by scaffolding and gapfilling. https://github.com/sanger-pathogens/assembly_improvement.

Wellcome Sanger Institute. Pathogen Informatics. https://github.com/sanger-pathogens/vr-codebase.

Wellcome Sanger Institute. SMALT: A mapper for DNA sequencing reads. http://sourceforge.net/projects/smalt/.

Wilson DN. 2014. Ribosome-targeting antibiotics and mechanisms of bacterial resistance. Nat Rev Microbiol 12: 35–48.

Young BC, Earle SG, Soeng S, Sar P, Kumar V, Hor S, Sar V, Bousfield R, Sanderson ND, Barker L, et al. 2019. Panton-Valentine leucocidin is the key determinant of Staphylococcus aureus pyomyositis in a bacterial GWAS. Elife 8. http://dx.doi.org/10.7554/eLife.42486.

Zaman S, Fitzpatrick M, Lindahl L, Zengel J. 2007. Novel mutations in ribosomal proteins L4 and L22 that confer erythromycin resistance in Escherichia coli. Mol Microbiol 66: 1039–1050.

Zerbino DR, Birney E. 2008. Velvet: algorithms for de novo short read assembly using de Bruijn graphs. Genome Res 18: 821–829.

