## Supplementary figures and images for "Contrasting approaches to genome-wide association studies impact the detection of resistance mechanisms in *Staphylococcus aureus*"

### Supplemental Fig. S1

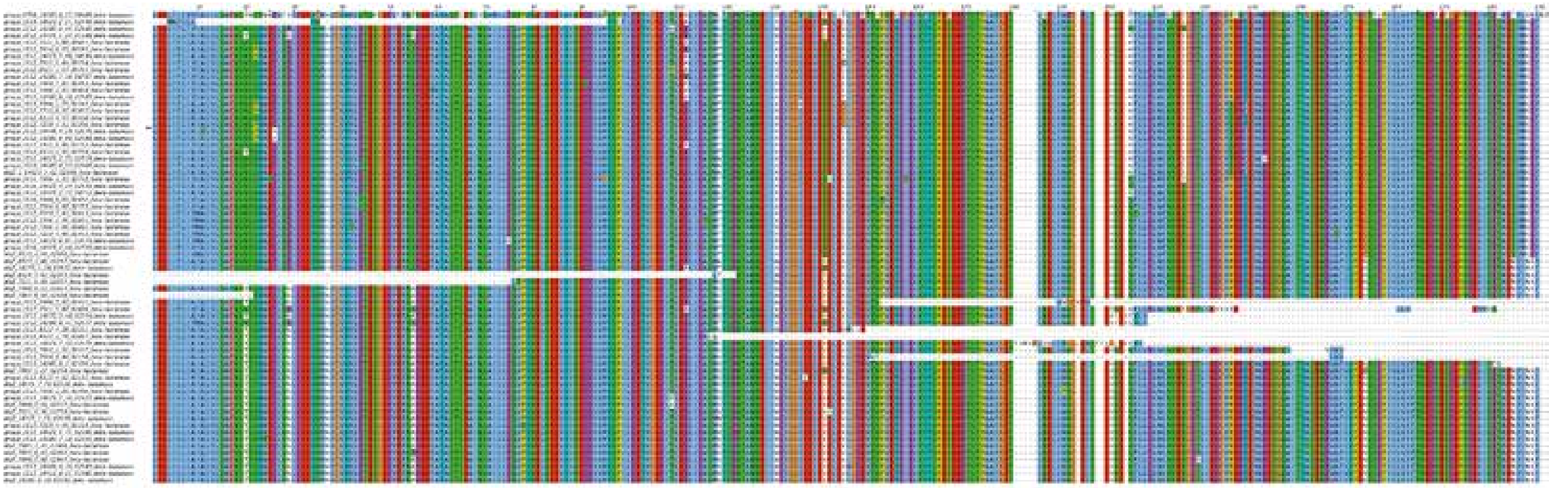

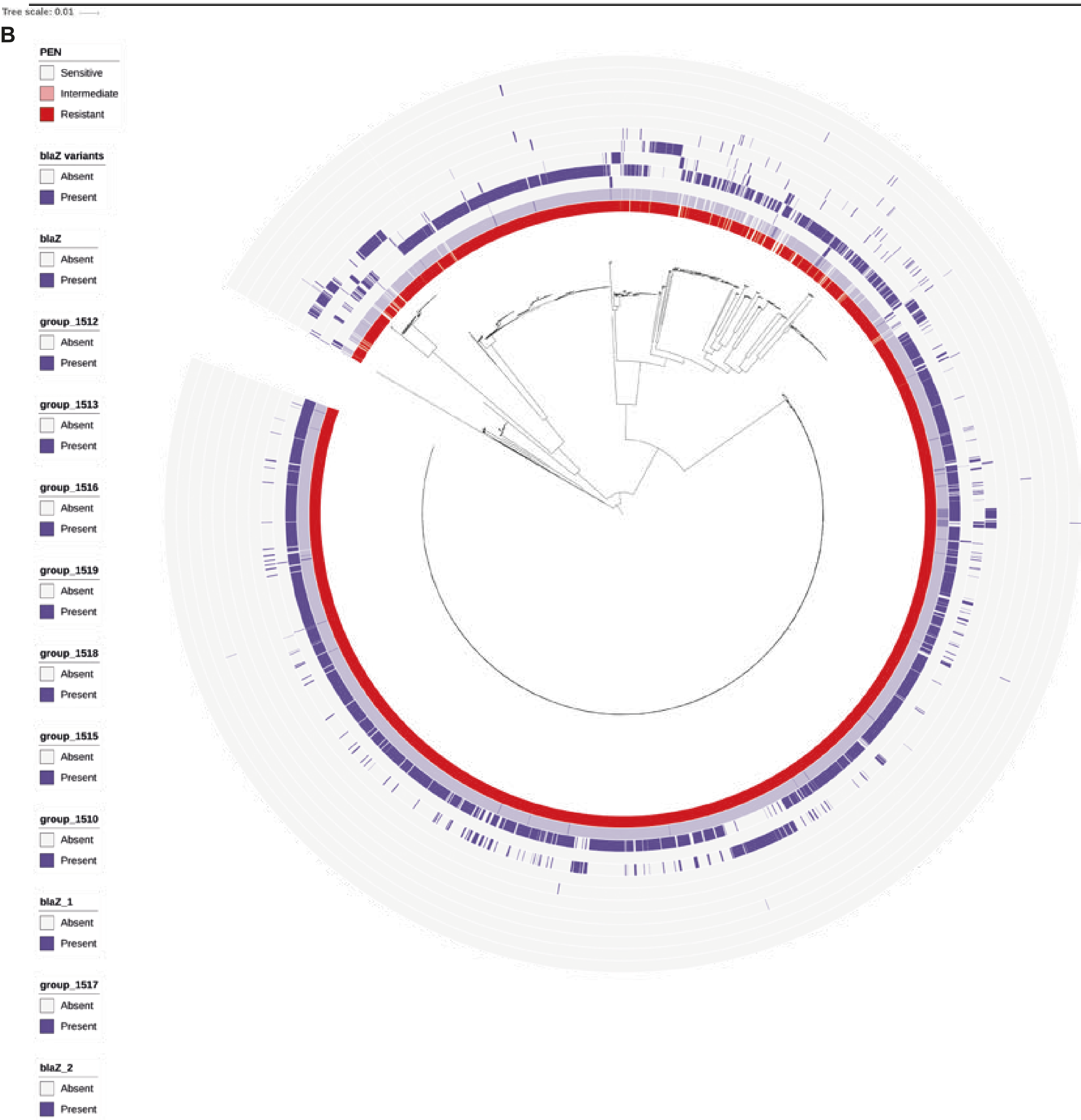

### Supplemental Fig. S2

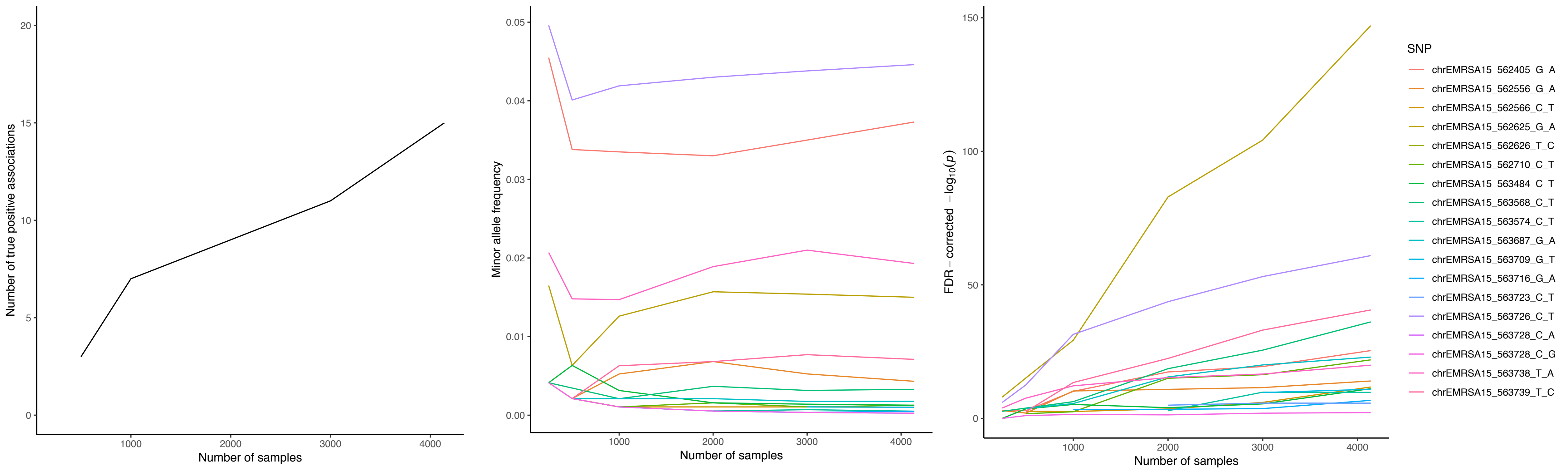

### Supplemental Fig. S4

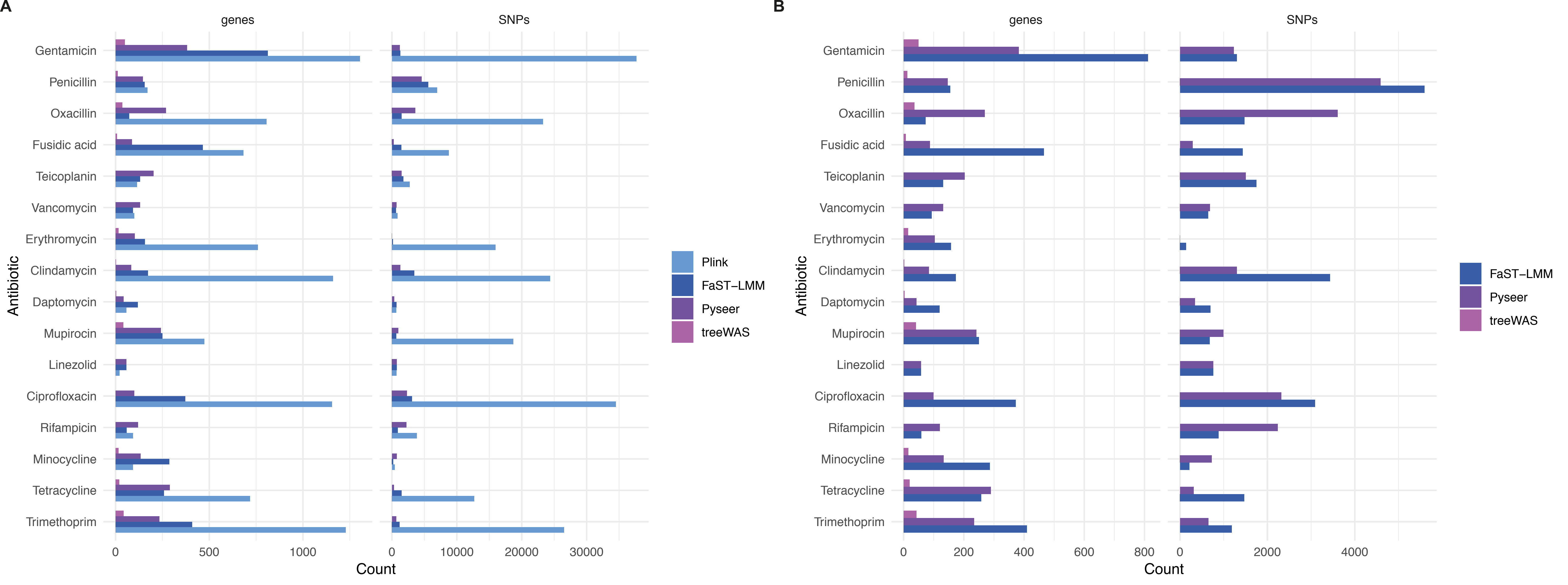
