## Supplemental Fig. S3 for "Contrasting approaches to genome-wide association studies impact the detection of resistance mechanisms in *Staphylococcus aureus*"

```
                  1
ileS_1            MDYKETLLMP KTDFPMRGGL PNKEPQIQEK WDAEDQYHKA LEKNKGNETF ILHDGPPYAN GNLHMGHALN KILKDFIVRY KTMQGFYAPY VPGWDTHGLP
ileS-2            LTKKYL---- ---------- -NTQNEISAF WNTQKIFKKS IDNRKGQESF VFYDGPPTAN GLPHAGHVLG RVIKDLVARL KTMQGFYVER KAGWDTHGLP
ileS-3_8140_1#33  LTHKKL---- ---------- -NTQHEISEF WNANHIFKKS MENRRDNEKF VFYDGPPTAN GLPHAGHVLG RVIKDLVSRF KTMQGFYVER KAGWDTHGLP

                101
ileS_1            IEQALTKK-G V----DRKKM STAEFREKCK EFALEQIELQ KKDFRR-LGV RGDFNDPYIT LKPEYEAAQI RIFGEMADKG LIYKGKKPVY WSPSSESSLA
ileS-2            VELEVEKKIG IKGKQDIEKY GIENFINECK KSVFN-YEKE WRDFSKDLGY WVDMDSPYIT LENNYIESVW NILSTFHKKG LLYKGHKVTP YCTHDQTALS
ileS-3_8140_1#33  VELEVEKKIG IEGKKDIENY GIENFINECK NSVFN-YAKE WRDFSNALGY WVDMDNPYIT LDNNYIESVW NILSTFHKQG LLYKGHKVTP YCTHDQTALS

                201
ileS_1            EAEIE--YHD KRSASIYVAF NVKDDKGVVD ADAKFIIWTT TPWTIPSNVA ITVHPELKYG QYNVNGEKYI IAEALSDAVA EALDWDKASI KLEKEYTGKE
ileS-2            SHEVAQGYKN VKDLSAVVKF QLTNSK---- -DTYFLSWTT TPWTLPANVA LAINKDLNYS KIRVENEYYI LATDLINSII ------TEKY EIIDTFSGSN
ileS-3_8140_1#33  SHEVAQGYKN VKDLSAIVKF QQKNDI---- -NTFFLSWTT TPWTLPANVA LAINKNMNYS KIQIENSYYI LATDLIDSII ------TEEY TFIETFSGKD

                301
ileS_1            LEYVVAQHPF ----LDRESL VINGDHVTTD AGTGCVHTAP GHGEDDYIVG QKYELPVISP IDDKGVFTEE GGQFEGMFYD KANKAVTDLL TEKGALLKLD
ileS-2            LINLKYIPPF ESDGLVNAYY VVDGEFVTNS EGTGIVHIAP AHGEDDYQLV LERDLDFLNV ITREGVYNDR FPELVGNKAK NSDIEIIKLL SKKQLLYKKQ
ileS-3_8140_1#33  LINLEYVPPF KNDNLTNAYY VVDAEFVTNS EGTGIVHIAP AHGEDDYKLA SENNLEFLNV ITIEGIYNYN FLELEGKKAK DSDIEIIKLL SKKKLLYKKQ

                401
ileS_1            FITHSYPHDW RTKKPVIFRA TPQWFASISK VRQDILDAIE NTN-FKVNWG KTRIYNMVRD RGEWVISRQR VWGVPLPVFY AENGEIIMTK ETVNHV-ADL
ileS-2            KYEHNYPHCW RCGNPLIYYA MEGWFIKTTN FKNEIINNNN NIEWFPSHIK EGRMGNFLEN MVDWNIGRNR YWGTPLNVWI CNDCNHEYAP SSIKDLQNNS
ileS-3_8140_1#33  KYEHSYPHCW RCGNPLIYYA MEGWFIKTTN FKEEIIKNNK NIDWFPSHIK DGRMGNFLEN MVDWNIGRNR YWGTPLNVWT CNDCNYEFSP NSIEDLRTNS

                501
ileS_1            FAEHGSNIWF EREAKDLLPE GFTHPGSPNG TFTKETDIMD VWFDSGS--- -SHRGVLETR PELS--FPAD MYLEGSDQYR GWFNSSITTS VATRGVSPYK
ileS-2            INKIDEDIEL HRPYVDNIT- -LSCPK-CNG KMSRVEEVID VWFDSGSMPF AQHHYPFDNQ KIFNQHFPAD FIAEGVDQTR GWFYSLLVIS TILKGKSSYK
ileS-3_8140_1#33  LNEVPKDIEL HRPYVDDII- -CHCPK-CNG KMTREEEVID VWFDSGSMPF AQHHYPFDDK KDFKQHFPAD FIAEGVDQTR GWFYSLLVIS TILKGQSSYK

                601
ileS_1            FLLSHGFVMD GEGKKMSKSL GNVIVPDQVV KQKGADIARL W--VSSTDYL ADVRISDEIL KQTSDVY-RK IRNTLRF--M LGNINDFNPD TDSIPESELL
ileS-2            RALSLGHILD SNGKKMSKSK GNVINPTELI NKYGADSLR- WALISDSAPW NNKRFSENIV AQTKSKFIDT LDNIYKFYNM YNKIDHYNPN NEITKSRNTL
ileS-3_8140_1#33  RAFSLGHILD SNGKKMSKSK GNVINPSDLI NKYGADSLR- WALISDSAPW NNKRFSESIV AQTKSKFIET LDNIYKFYNM YNEIDKFKPQ SQY--SENLL

                701
ileS_1            EVDRYLLNRL REFTASTINN YENFDYLNIY QEVQNFINVE LSNFYLDYGK DILYIEQRDS HIRRSMQTVL YQILVDMTKL LAPILVHTAE EVWSHTPHVK
ileS-2            --DNWALSRL NTLIKESNIY VNNYDFTSAA RLINEYTNT- ISNWYIRRSR GRFW-EQGIS NDKKDAYNTL YEILTTLSRL VAPFVPFISE KIHY---NLT
ileS-3_8140_1#33  --DKWALSRL NTLIKETNTH INNYDFTSAS RLINDFTSL- ISNWYIRRSR SRFW-EEGLS DDKINAYNTL YEILINLSKL IAPFTPFISE KIHY---NLT

                801
ileS_1            EESVHLADMP KVVE--VDQA LLDKWRTFMN LRDDVNRALE TARNEKVIGK SLEAKVTIAS NDKFNASEFL TSFDALHQLF IVSQVKVVDK LDDQATAYE-
ileS-2            GKSVHLQDYP QYKESFINQA LEDEMHTVIK I---VELSRQ ARKNADLKIK QPLSKMVIKP NSQLNLSFLP NYYSIIKDEL NIKNIELTDN INDYIT-YEL
ileS-3_8140_1#33  GESVHLTNYP QHNDSLINRN LEDEMYTVIK I---VELARQ ARKNANLKVK QPLSELIIKS NKQLNLNFLS NYNSIIKDEL NIKNVKIVSN INDYIS-YDI

                901
ileS_1            ---------- ---------- ---------- ---------- ---------H GDIVIEH--- -----ADGEK CERCWN---- ---------- ----------
ileS-2            KLNFSSVGPK LGNKTKNIQT LIDSLSEYDK KSLIESNNFK SLSSDAELTK DDFIIKTLPK DSYQLSEDND CVILLDKNLS PELIREGHAR ELIRLIQQLR
ileS-3_8140_1#33  NLNFSSAGPK LGNKVKEVKT SLNSLSEKDK KNLVESKDFQ KLSSDILLTD EDFVIKVLPI ENYQLSDDSD YSVLLNKKLS SELIHEGNAR ELIRLIQQLR

               1001
ileS_1            ---------- ---------- ---------- -YSED----- --LGAVDELT HLCPRCQQVV K-SLV*-
ileS-2            KKKNLPINQR IDIYIGVTGE LLESIKTNKN MFKENLLIKN IHLNVIDEYE NTIHFNNKEI KISLLY*
ileS-3_8140_1#33  KKKDLPINQR IDLHIGVKGD LLDSINSNKG MFKENLLINN FSLNTLDEYE DIISLNNKEI KISLLY*
```
