## Supplemental Table S1 for "Contrasting approaches to genome-wide association studies impact the detection of resistance mechanisms in *Staphylococcus aureus*"

Table S1. Source of isolates and method of susceptibility testing.

| Collection | Isolate number | Source | Date | Sample type | Susceptibility testing |
| --- | --- | --- | --- | --- | --- |
| Dataset 1  (n=1,818) | 1,013 MRSA | Submitted to the BSAC bacteraemia resistance surveillance programme by 47 laboratories in the United Kingdom and Republic of Ireland (Reuter et al. 2016). | 2001-2010 | Blood | Agar dilution method |
|  | 538  (134 MRSA, 404 MSSA) | Cambridge University Hospitals NHS Foundation Trust, Cambridge (CUH) United Kingdom (Donker et al. 2017) | 2006-2012 | Blood | Automated instrument (Vitek2) |
|  | 269 MRSA | 9 hospitals in the East of England (Donker et al. 2017) | 1998-2012 | Blood | Automated instrument (Vitek2) |
| Dataset 2  (n=2,322) | 2,322 MRSA | At least one isolate from all MRSA positive cases identified by the Public Health England Clinical Microbiology and Public Health Laboratory, CUH over 12 months. Samples were received from four hospitals and 75 primary care practices. (Coll et al. 2017) | 2012-2013 | Blood (65)  Other clinical sample type (638)  Screening swab (1,619) | Automated instrument (Vitek2) |

References

Coll F, Harrison EM, Toleman MS, Reuter S, Blane B, Raven KE, Palmer B, Kappeler ARM, Brown NM, Torok ME et al. 2017. Extensive cryptic transmission of MRSA revealed by population-based longitudinal genomic surveillance. *Science Translational Medicin* **accepted**.

Donker T, Reuter S, Scriberas J, Reynolds R, Brown NM, Torok ME, James R, Network EoEMR, Aanensen DM, Bentley SD et al. 2017. Population genetic structuring of methicillin-resistant Staphylococcus aureus clone EMRSA-15 within UK reflects patient referral patterns. *Microbial Genomics* **accepted**.

Reuter S, Torok ME, Holden MT, Reynolds R, Raven KE, Blane B, Donker T, Bentley SD, Aanensen DM, Grundmann H et al. 2016. Building a genomic framework for prospective MRSA surveillance in the United Kingdom and the Republic of Ireland. *Genome research* **26**(2): 263-270.
